# Afferent connections of cytoarchitectural area 6M and surrounding cortex in the marmoset: putative homologues of the supplementary and pre-supplementary motor areas

**DOI:** 10.1101/2021.05.21.445121

**Authors:** Sophia Bakola, Kathleen J. Burman, Sylwia Bednarek, Jonathan M. Chan, Natalia Jermakow, Katrina H. Worthy, Piotr Majka, Marcello G.P. Rosa

**Affiliations:** Department of Physiology and Biomedicine Discovery Institute, Monash University, Clayton, VIC 3800, Australia; ARC Centre of Excellence for Integrative Brain Function, Monash University Node, Monash University, Clayton, VIC 3800, Australia; Laboratory of Neuroinformatics, Nencki Institute of Experimental Biology of the Polish Academy of Sciences, 02-093 Warsaw, Poland

**Author notes:** These authors contributed equally to this work and are listed here in alphabetical order (Sophia Bakola, Kathleen J. Burman). Joint senior authors (Piotr Majka, Marcello G.P. Rosa).

## Abstract

Cortical projections to the caudomedial frontal cortex were studied using retrograde tracers in marmosets. We tested the hypothesis that cytoarchitectural area 6M includes homologues of the supplementary and pre-supplementary motor areas (SMA and preSMA) of other primates. We found that, irrespective of the injection sites’ location within 6M, over half of the labeled neurons were located in motor and premotor areas. Other connections originated in prefrontal area 8b, ventral anterior and posterior cingulate areas, somatosensory areas (3a and 1-2), and areas on the rostral aspect of the dorsal posterior parietal cortex. Although the origin of afferents was similar, injections in rostral 6M received higher percentages of prefrontal afferents, and fewer somatosensory afferents, compared to caudal injections, compatible with differentiation into SMA and preSMA. Injections rostral to 6M (area 8b) revealed a very different set of connections, with increased emphasis in prefrontal and posterior cingulate afferents, and fewer parietal afferents. The connections of 6M were also quantitatively different from those of M1, dorsal premotor areas, and cingulate motor area 24d. These results show that the cortical motor control circuit is conserved in simian primates, indicating that marmosets can be valuable models for studying movement planning and control.

## Introduction

The primate cortical motor planning and control network represents a significant elaboration of the basic plan shared with other mammals, notably in terms of increased number of anatomical and functional subdivisions in the caudal frontal (premotor) and posterior parietal regions (Kaas 2012; Kaas and Stepniewska 2016). Although there are indications that the main components of this network are conserved across the main taxonomic groups of primates (Geyer et al. 2000; Bakola et al. 2015; Borra and Lupino 2019), we are still far from understanding the extent of the variability between species, which is essential for the informed translation of experimental results from preclinical models to humans.

The present paper is part of a series of studies focused on the network of neuronal connections underlying motor control in the common marmoset (*Callithrix jacchus*), a New World monkey species that is becoming increasingly used in neuroscience. Marmosets are among the smallest primates, and offer advantages relative to the more widely used macaque monkey in terms of shorter developmental time (Burman et al. 2007, Sawiak et al. 2018), simpler brain morphology, which facilitates the integration of data across individuals (Majka et al. 2016, 2020) and fewer biosafety concerns (Mansfield 2003). Conversely, certain aspects of marmoset brain circuitry may reflect differences in motor behavior, such as the capacity to perform independent finger movements (Padberg et al. 2007). Previous studies have mapped the afferent connections of the primary motor, dorsal and ventral premotor areas in this species (Burman et al. 2014a, b, 2015), but there have been no studies of the putative premotor areas located near the dorsal midline. Given the rapidly developing use of marmosets in studies of motor control (Bakola et al. 2015; Roy et al. 2016; Tia et al. 2017; Walker et al. 2017; Ebina et al. 2018, 2019; Pomberger et al. 2019), improved knowledge of the pattern of interconnections between cortical areas is likely to lead to better models to understand the mechanistic basis of neuronal interactions leading to the planning and execution of movement, as well as the circuit basis of differences in motor behavior.

In primates, the premotor cortex in the dorsomedial convexity of the hemisphere is usually subdivided into supplementary (SMA, or F3) and pre-supplementary (preSMA, or F6) motor areas, which are both considered subdivisions of cytoarchitectural area 6 (Barbas and Pandya 1987; Luppino et al. 1991; Matelli et al. 1991; Tanji and Shima 1994; Akkal et al. 2007). In macaques, the preSMA has distinct anatomical features, including more numerous connections with prefrontal and non-motor areas (Bates and Goldman-Rakic 1993; Luppino et al. 1993; Lu et al. 1994). Like SMA, the preSMA engages in motor acts but becomes activated at more abstract levels, such as during the temporal coding of sequential behavior (Picard and Strick 1996; Tanji 2001). Presently, there is no clear evidence of a similar subdivision in the marmoset, in which a single cytoarchitectural field (area 6M) has been described in the corresponding location (Burman et al. 2008; Paxinos et al. 2012). However, the nature of the anatomical transitions between area 6M and the areas it adjoins has not been characterized. To address these questions, we report on the results of experiments involving retrograde tracer injections in marmosets and computational reconstruction of connection patterns.

## Materials and methods

### Animals and tracer injections

The experiments conformed to the Australian Code of Practice for the Care and Use of Animals for Scientific Purposes and were approved by the Monash University Animal Experimentation Ethics Committee. Nine young adult marmoset monkeys (*Callithrix jacchus*, the common marmoset) received 12 injections of neuronal tracers in the frontal cortex near the midline, within and around the expected location of area 6M according to the Paxinos et al. (2012) atlas. To reduce the number of experimental animals, they each received multiple distinct tracer injections in other locations, as part of past and ongoing projects in this laboratory. The actual locations of the injections, based on histological reconstruction, are summarized in Table 1. Figure 1 illustrates the injection site locations following registration to the stereotaxic space of the Paxinos et al. (2012) marmoset brain atlas, and Figure 2 shows their extents relative to the underlying histology. The data obtained following 10 fluorescent tracer injections were processed using a computational pipeline (Majka et al. 2016, 2020), thus being also available in full through the Marmoset Brain Connectivity Atlas web site (www.marmosetbrain.org, RRID: SCR_015964). This includes one injection (case 9, CJ83-DY) from a previous study (Reser et al. 2013), which has been fully reanalyzed. Two injections, which used the tracer biotinylated dextran amine (BDA), were placed in hemispheres contralateral to those receiving fluorescent tracers and only processed using manual reconstruction techniques (see Burman et al. 2014a, b). Quantitative comparisons with adjacent motor areas (M1, 6DC and 6DR) were based on previously published results (Burman et al. 2014a, b; also available through www.marmosetbrain.org).

**Table 1.** Summary of experimental cases.

| Case | Designation | Sex | Age<br>(years,<br>months) | Weight<br>(g) | Hemisphere | Injection<br>site | Survival<br>(days) | Amount,<br>concentration | Number of<br>labeled<br>cells<sup>b</sup> | Layers<br>involved<sup>c</sup> |
| --- | --- | --- | --- | --- | --- | --- | --- | --- | --- | --- |
| 1 | CJ94-FB<sup>a</sup> | F | 3y 2m | 397 | R | 6M | 14 | 0.3 µl, 1.5% | 4830 | 1-6 |
| 2 | CJ115-BDA | F | 2y 10m | 387 | L | 6M | 17 | 0.6 µl, 11% | 1734 | 1-5 |
| 3 | CJ112-BDA | M | 3y 1m | 377 | R | 6M | 17 | 0.6 µl, 11% | 4430 | 1-6 |
| 4 | CJ113-FE | M | 3y 1m | 423 | R | 6M | 17 | 0.3 µl, 15% | 2435 | 1-5 |
| 5 | CJ163-FB | F | 1y 6m | 324 | L | 6M | 17 | 0.3 µl, 1.5% | 22445 | 1-6 |
| 6 | CJ170-DY | F | 2y 3m | 326 | R | 6M | 17 | 0.3 µl, 1.5% | 1967 | 1-5 |
| 7 | CJ164-CTBg | F | 1y 5m | 304 | R | 6M | 17 | 0.3 µl, 1% | 2584 | 1-5 |
| 8 | CJ164-FB | F | 1y 5m | 304 | R | 8b | 14 | 0.3 µl, 1.5% | 25529 | 1-4 |
| 9 | CJ83-DY | F | 2y 5m | 329 | R | 8b | 14 | 0.4 µl, 1.5% | 8574 | 1-6 |
| 10 | CJ163-DY | F | 1y 6m | 324 | L | 8b | 17 | 0.5 µl, 1.5% | 12807 | 1-6 |
| 11 | CJ164-DY | F | 1y 5m | 304 | R | 8b | 17 | 0.3 µl, 1.5% | 24282 | 1-4 |
| 12 | CJ167-CTBg | F | 2y 2m | 390 | R | 24d | 16 | 0.3 µl, 1% | 9093 | 1-4 |

<sup>a</sup>Tracer abbreviations: CTBg, cholera toxin subunit B conjugated with Alexa 488; DY, diamidino yellow dihydrochloride; FE, fluoroemerald; FB, fast blue; BDA, biotinylated dextran amine.
<sup>b</sup>Extrinsic ipsilateral projections.
<sup>c</sup>Range of layers that were at least partially involved, across all sections containing the injection site

**Figure 1.**
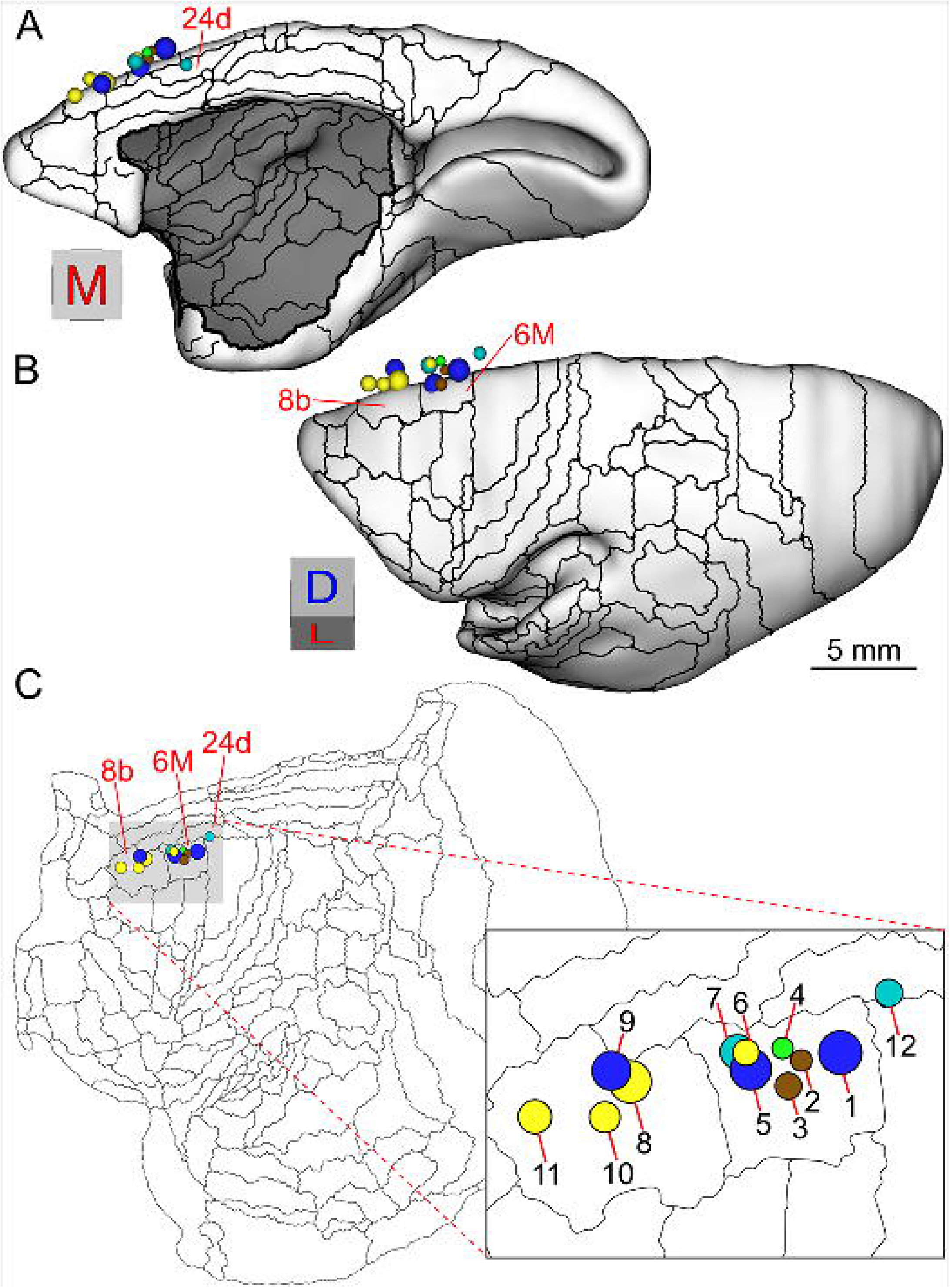
Summary of the locations of 12 injection sites, relative to the borders of marmoset frontal areas. **A** and **B** are medial and dorsolateral views of a computerized reconstruction of the left hemisphere of the marmoset brain, at the level of the cortical mid-thickness. The images represent the template generated by Majka et al. (2016) on the basis of the coronal plates provided by Paxinos et al. (2012). Thin lines represent the boundaries of areas, and the orientation cubes next to each brain image indicate orientation (M-medial, D-dorsal, L-lateral). The areas which received injections in the present study (6M, 8b, 24d) are indicated. The locations of the spheres correspond to the centers of mass of the injection sites, and the volumes of the spheres are proportional to the histologically determined injection core zones. Dark blue indicates FB injections, yellow indicates DY injections, brown indicates BDA injections, teal indicates CTB injections, and light green indicates a FE injection. The actual shape of the injection sites varied, with most corresponding to elongated tracer deposits (see Fig. 2). In **C**, the same information is shown projected radially onto a flat map of the marmoset cortex (frontal to the left, cingulate to the top), with the injection site case numbers identified according to Table 1.

**Figure 2.**
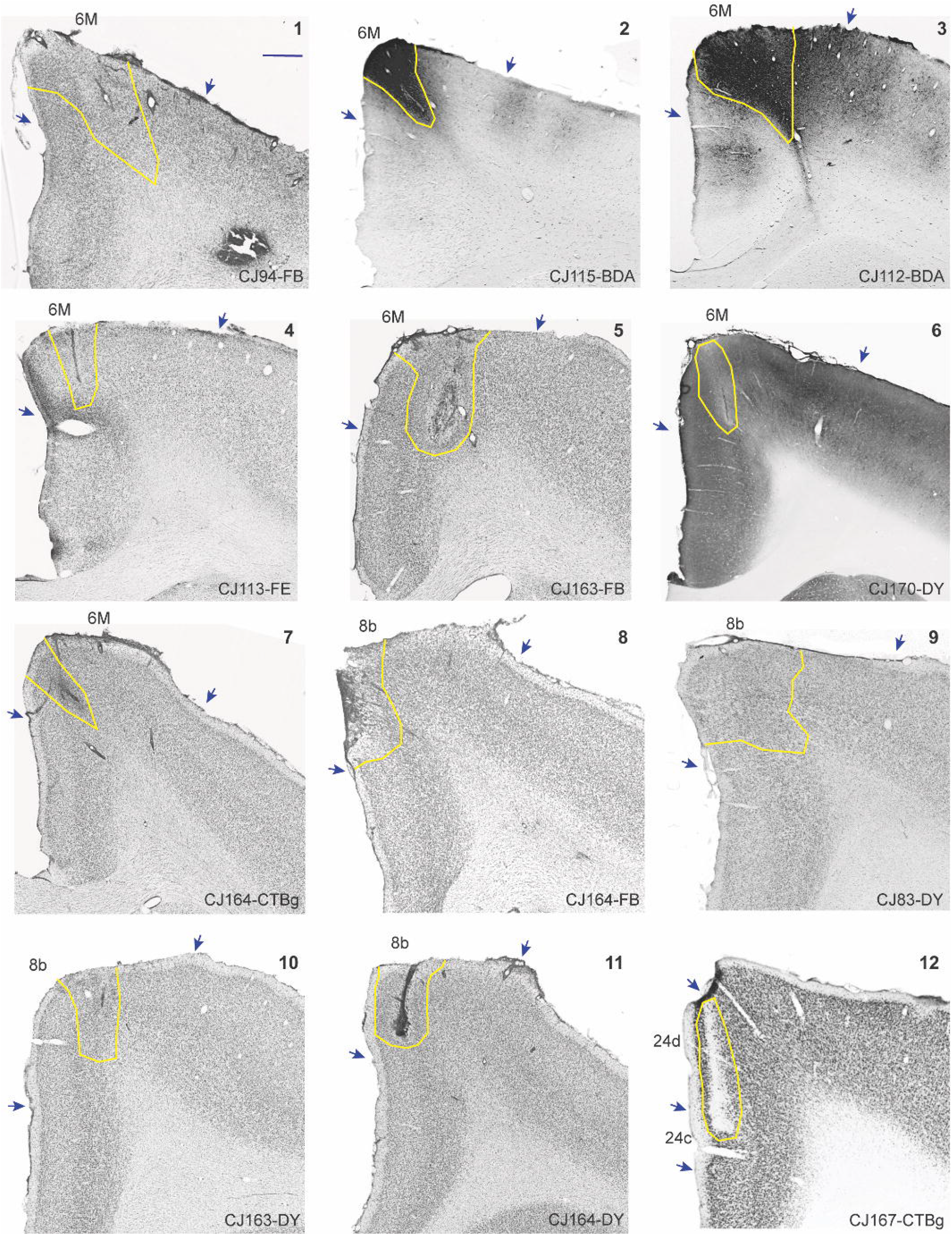
Coronal sections through the centers of the injection sites. The panels are numbered 1-12 (top right) according to Table 1 and Figure 1. Each panel illustrates the brain’s dorsomedial convexity, with the extent of the injection site (core + halo, outlined under the microscope), indicated in yellow, and the borders of the area that received the injection are indicated by arrows. For cases 1, 4, 5, and 7-11 the illustrated sections were stained for Nissl bodies using cresyl violet, for cases 2 and 3 they were reacted to reveal the distribution of BDA, for case 6 it was reacted for cytochrome oxidase, and for case 12 it was immunohistochemically stained for NeuN. Scale bar (top left panel) = 1 mm. Full views of the injection sites are available through www.marmosetbrain.org.

Surgical procedures have been reported in detail in previous papers (Reser et al. 2013; Burman et al. 2014a, 2014b; Burman et al. 2015). Briefly, intramuscular (i.m.) injections of atropine (0.2 mg/kg) and diazepam (2 mg/kg) were administered as pre-medication; 30 min later, anesthesia was achieved with alfaxalone (Alfaxan, 10 mg/kg, i.m.), with supplemental doses of the drug administered as needed during surgery to maintain areflexia. Body temperature, heart rate and blood oxygenation were continuously monitored during procedures. Dexamethasone (0.3 mg/kg, i.m.) and amoxicillin (50 mg/kg, i.m.) were also administered before the animals were placed in the stereotaxic frame. Selection of injection sites was guided by stereotaxic coordinates, estimated from previous studies using the interaural level as reference (Paxinos et al. 2012). Assignment of injection sites to areas was done based on *post-mortem* histological analysis, as detailed below.

To identify projections, we used BDA (10 kDa, Invitrogen-Molecular Probes) and four fluorescent tracers: fluoroemerald (FE; dextran-conjugated fluorescein 10 kDa; Life Technologies), cholera toxin subunit B conjugated with Alexa 488 (CTBgr, Invitrogen-Molecular Probes), diamidino yellow dihydrochloride (DY, Polysciences) and fast blue (FB, Polysciences). BDA and FE resulted in bidirectional transport, but only retrograde label (which can be readily quantified) is reported here. Tracers were injected slowly, in steps of 40 nl over 15-20 min, using a microsyringe fitted with a glass micropipette. The animals were monitored for several hours postoperatively and administered analgesics (Temgesic 0.01 mg/kg, i.m., once, and Carprofen 4 mg/kg, subcutaneously for 2–3 days after surgery).

After a survival period between 14 and 17 days (Table 1), the animals were re-anaesthetized as above and, once unconscious, administered a lethal dose of sodium pentobarbitone (100 mg/kg, intraperitoneally). Once respiratory arrest was observed, they were perfused through the heart, with heparinized saline, followed by 4% paraformaldehyde in 0.1 M phosphate-buffered saline (pH 7.4). The brains were removed from the skull, post-fixed for up to 24 h, and cryoprotected by immersion in ascending concentrations of sucrose (10–30% in buffered paraformaldehyde). Using a cryostat, 40 μm thick sections were cut in the coronal plane and collected into five consecutive series. All sections from one series were mounted unstained for the examination of fluorescent labeling. In animals CJ112 and CJ115, another series was processed for BDA histochemistry using an ABC kit (Vector Laboratories, Burlingame, CA) and 3,3**’**-diaminobenzidine with nickel salt as an enhancer (Reser et al. 2017). Sections from the remaining series were stained for Nissl substance, cytochrome oxidase and myelin, using publicly available protocols optimized for non-human primate tissue (Worthy and Burman 2017; Worthy and Rosa 2017; Worthy et al. 2017; available at http://www.marmosetbrain.org/reference). All sections were air-dried overnight and coverslipped with di-n-butyl phthalate xylene (DPX).

### Data analysis

The locations of cells labeled by the different tracers were charted using a digitizing system (MD3 digitizer and MDPlot software, Accustage) interfaced to a Zeiss Axioskop 2 microscope. This analysis was conducted in every fifth section (200 µm separation, most cases), except for one case (CJ94-FB), in which every tenth section was examined.

Attribution of injection sites and labeled neurons to architectonic fields was determined in histological material, using as a guide the designations in our previous papers (Reser et al. 2013; Burman et al. 2014a, 2014b; Burman et al. 2015) and those in the marmoset brain atlas (Paxinos et al. 2012). As described previously (Burman et al. 2006; Burman et al. 2008) and illustrated in Figure 3, area 6M can be distinguished from neighboring areas based on cyto- and myeloarchitectural criteria. In myelin-stained sections (Fig. 3B-C, left) area 6M is characterized by relatively dense myelination, thick radial bundles of fibers emerging from the white matter spanning the lower layers, and poor separation between the bands of Baillarger. The rostral and lateral transitions, respectively to area 8b (Fig. 3A, left) and to area 6DR (dorsorostral premotor area; Fig. 3B, left), laterally, are accompanied by a reduction in the overall myelination, and the appearance of a slightly more lightly myelinated gap at the approximate level of layer 4, which results in an incipient separation between the inner and outer bands of Baillarger. Transitions to both area 6DC (dorsocaudal premotor area), laterally, and to the medial part of M1 (cytoarchitectural area 4a), caudally, are characterized by an increase in myelination, particularly at the level of layer 3 (Fig. 3C-D, left).

**Figure 3.**
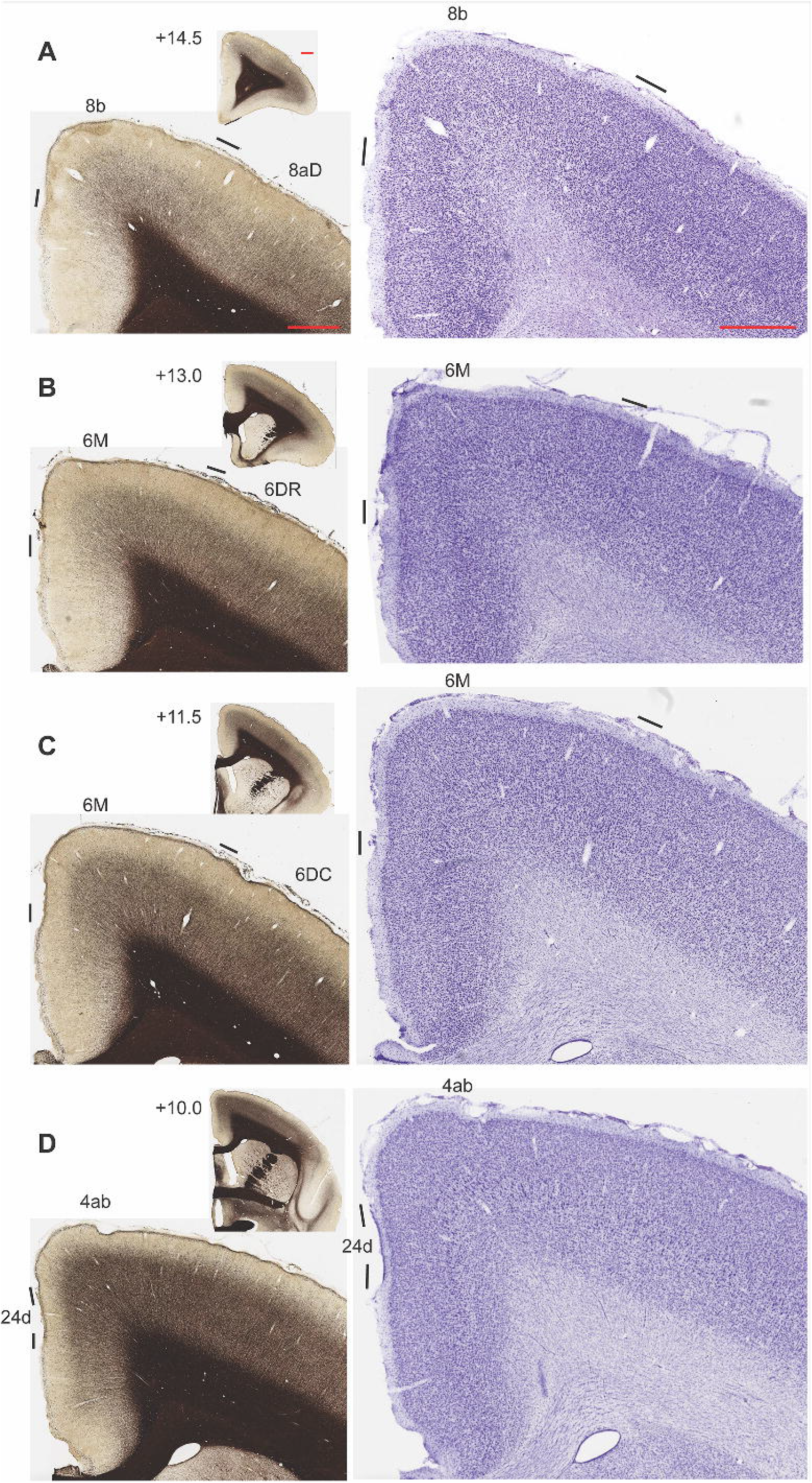
Histological characteristics of the marmoset dorsomedial frontal cortex areas that are part of this study. Panels **A-D** illustrate adjacent sections stained for myelin (left) and Nissl bodies (right). Histological borders between areas are indicated as transition zones by black bars above layer 1. The anteroposterior stereotaxic level of each section is indicated in the inserts adjacent to the myelin-stained sections. For a description of the histological patterns and criteria for transitions, see “Materials and Methods”. Scale bars (1 mm) are shown as red lines in panel A.

Changes in cytoarchitecture accompany these myeloarchitectural transitions. Area 6M is distinguished from M1, 6DR and 6DC by the presence of a slight increase in cell density in the middle layers, including the expected level of layer 4 (Fig. 3B-C, right), which is not apparent in the latter areas. The rostral transition to area 8b is accompanied by a better differentiation of layer 4, and a reduction in cellular density in the infragranular layers (Fig. 3A, right). In addition, the transitions to 6DC and M1 reveal larger neurons in layer 5 (particularly in the latter), and lower neuronal density in the infragranular layers (Fig. 3C-D, right; see also Atapour et al. 2019). Finally, the borders of area 6M with subdivisions of the putative cingulate motor cortex (areas 24c and 24d) were also obvious. Among other characteristics, the transition to area 24c coincided with a drop in myelination (Fig. 3B-C, left), and that to area 24d by the occurrence of larger neurons in layer 5 in this area (Fig. 3D, right), which were however smaller than those found in M1. As detailed below, the connections pattern also suggested a differentiation between the rostral and caudal parts of area 6M. Although we could not detect a reliable transition using the present histological methods, there was a tendency for the caudal part of area 6M to be more myelinated compared to the rostral parts (compare Fig. 3B and C, left).

Data from each of the cases with injections of fluorescent tracers were computationally registered onto a volumetric template based on the Paxinos et al. (2012) brain atlas, and visualized in two-dimensional reconstructions of the template cortical surface (see Majka et al. 2016, 2019). To prepare the reconstructions, results from each hemisphere were exported from the MDplot software in text format (MDO), transformed into scalable vector graphics (SVG) and aligned to the nearest scanned images of histological sections; alignment involved simple manipulations, usually rotation, translation and/or scaling of experimental sections (Majka et al. 2016). Finally, co-registration of single hemispheres with the template brain was achieved through affine transformation and warping, based on prominent cortical landmarks such as the lips and fundus of the calcarine and lateral sulcus. The reconstruction procedure also used histological borders visualized in each individual animal as landmarks, and was followed by expert evaluation to ensure accuracy in all regions containing labeled cells (Majka et al. 2020). The hemispheres injected with BDA, which were not registered to the template due to incompatibility with the computational pipeline, were analyzed section by section using manual annotation in relation to with adjacent myelin, cytochrome oxidase and Nissl-stained sections.

## Results

We report on the results of 12 retrograde tracer injections in the caudal frontal cortex of 9 marmosets. The injections were placed near the dorsal midline, in some cases also involving the medial wall of the hemisphere (Fig. 1). Reconstruction of the injection sites (Fig. 2) showed that layers 1-4 were always included, whereas infragranular layers were involved in most, but not all cases (Table 1). Histological analyses demonstrated that 7 injections were centered within cortical area 6M, 4 in area 8b, and 1 in area 24d (Table 1). Table 2 summarizes the percentages of labeled neurons forming extrinsic connections to area 6M, and Table 3 provides similar information for the injections in areas 8b and 24d. As described below, the analyses also involve quantitative comparisons with data reported by Burman et al. (2014a, b), including injections in areas 6DC, 6DR and M1.

**Table 2:** percentages of labelled neurons in different areas following injections in area 6M.

|  | **CJ94-**<br>**FB** | **CJ115-**<br>**BDA** | **CJ112-**<br>**BDA** | **CJ113-**<br>**FE** | **CJ163-**<br>**FB** | **CJ170-**<br>**DY** | **CJ164-**<br>**CTBg** |
| --- | --- | --- | --- | --- | --- | --- | --- |
| ***Granular frontal areas*** | ***1.13%*** | ***1.84%*** | ***0.84%*** | ***9.87%*** | ***12.27%*** | ***33.17%*** | ***27.62%*** |
| Frontal pole (area 10) | - | - | - | - | - | - | - |
| Dorsolateral prefrontal (areas 9, 46) | * | - | - | - | * | - | 0.34% |
| Ventrolateral prefrontal (areas 45, 47) | - | - | * | - | 0.60% | - | 0.41% |
| Orbitofrontal (areas 11, 13) | * | - | - | * | 0.20% | * | 2.14% |
| Area 8b | 0.82% | 1.79% | 0.72% | 9.63% | 8.86% | 31.85% | 16.18% |
| Area 8aD | * | * | * | * | 1.14% | 0.95% | 4.50% |
| area 8aV | 0.15% | - | - | * | 1.40% | 0.29% | 4.05% |
| ***Medial prefrontal and anterior cingulate areas*** | ***4.98%*** | ***4.28%*** | ***2.10%*** | ***2.43%*** | ***10.43%*** | ***1.81%*** | ***5.89%*** |
| Medial prefrontal (areas 14, 32V, 32) | - | - | - | - | * | - | - |
| Ventral anterior cingulate (area 24a) | * | 0.16% | * | - | 0.47% | 0.12% | - |
| Dorsal anterior cingulate (area 24b) | 4.96% | 4.12% | 2.05% | 2.43% | 9.90% | 1.69% | 5.89% |
| Perigenual (area 25) | - | - | - | - | * | - | - |
| ***Motor and premotor areas*** | ***80.05%*** | ***58.06%*** | ***73.86%*** | ***72.59%*** | ***61.89%*** | ***58.12%*** | ***61.72%*** |
| Primary motor (area 4a-c) | 39.18% | 17.53% | 21.22% | 8.54% | 4.41% | 1.03% | 10.23% |
| Medial premotor (area 6M) | i | i | i | i | i | i | i |
| Dorsocaudal premotor (area 6DC) | 7.60% | 14.70% | 27.72% | 20.13% | 24.06% | 20.92% | 16.01% |
| Dorsorostral premotor (area 6DR) | 0.42% | 7.33% | 4.24% | 18.32% | 11.10% | 6.04% | 12.16% |
| Area 8C | - | - | 0.61% | 0.39% | 0.45% | 3.37% | 0.21% |
| Ventral premotor (areas 6Va, 6Vb) | 5.23% | * | 2.42% | - | 3.10% | 3.99% | 0.31% |
| Cingulate motor (areas 24c, 24d) | 27.14% | 18.50% | 17.54% | 25.21% | 18.44% | 22.69% | 22.25% |
| Proisocortical motor (area ProM) | 0.48% | - | 0.11% | - | 0.32% | * | 0.55% |
| ***Somatosensory areas*** | ***5.87%*** | ***9.72%*** | ***9.11%*** | ***1.88%*** | ***1.46%*** | ***0.20%*** | ***0.16%*** |
| Area 3a | 2.41% | 0.60% | 2.17% | 0.16% | 0.40% | 0.16% | - |
| Area 3b | 0.17% | 1.09% | 1.47% | 0.23% | * | - | - |
| Area 1-2 | 1.35% | 5.97% | 3.50% | 1.10% | 0.13% | * | - |
| Lateral somatosensory (Areas S2, S2PV, S2PR) | 1.84% | 1.95% | 1.65% | 0.39% | 0.68% | - | * |
| Somatosensory insular (areas GI, ReI) | 0.10% | 0.11% | 0.32% | - | 0.23% | - | 0.13% |
| ***Posterior parietal areas*** | ***5.95%*** | ***24.04%*** | ***11.49%*** | ***9.08%*** | ***7.28%*** | ***5.25%*** | ***0.90%*** |
| Rostral posterior parietal (area PE) | 4.36% | 18.99% | 5.06% | 4.15% | 3.24% | 3.90% | * |
| Medial posterior parietal (areas PGM, 31, 23c) | 1.06% | 3.85% | 4.38% | 4.23% | 2.28% | 0.66% | 0.79% |
| Visuomotor association (areas V6a, PEC, MIP, VIP, LIP, AIP) | * | 0.81% | 0.81% | 0.31% | 0.35% | 0.12% | * |
| Rostral ventral parietal (areas PFG, PF) | 0.46% | 0.33% | 1.15% | 0.39% | 1.35% | 0.49% | * |
| Caudal ventral parietal (areas OPt, PG) | * | * | - | - | * | * | - |
| Temporoparietal transition (area TPt) | - | - | * | - | * | - | - |
| ***Posterior cingulate and retrosplenial areas*** | ***1.98%*** | ***1.52%*** | ***2.71%*** | ***2.58%*** | ***6.39%*** | ***1.39%*** | ***3.64%*** |
| Area 29 | * | - | * | * | 0.79% | 0.12% | * |
| Area 30 | * | 0.11% | * | - | 1.62% | 0.12% | 0.38% |
| Area 23a | 1.24% | 0.54% | 0.74% | 1.72% | 2.58% | 0.74% | 1.49% |
| Area 23b | 0.70% | 0.87% | 1.87% | 0.78% | 1.33% | 0.41% | 1.70% |
| Areas 23V and Prostriata | - | - | - | - | * | - | - |
| ***Orbital and temporal limbic areas*** | ***0.04%*** | ***-*** | ***-*** | ***-*** | ***0.20%*** | ***0.04%*** | ***0.07%*** |
| Gustatory cortex (area Gu) | - | - | - | - | * | - | * |
| Orbital and insular proisocortical areas (OPro, IPro, PaIM) | - | - | - | - | * | - | - |
| Orbital periallocortex (OPal) | - | - | - | - | * | - | - |
| Agranular and dysgranular insula | * | - | - | - | * | * | - |
| Piriform cortex | - | - | - | - | - | - | - |
| Entorhinal cortex | - | - | - | - | - | - | - |
| Perirhinal and temporal pole areas (areas 35, 36, TPro) | - | - | - | - | - | - | - |
| Parahippocampal (areas TH, TL, TF) | - | - | - | - | * | - | - |
| ***Temporal and occipital sensory areas*** | ***-*** | ***0.11%*** | ***-*** | ***-*** | ***0.08%*** | ***-*** | ***-*** |
| Auditory lateral belt (areas CL, ML, AL) | - | - | - | - | - | - | - |
| Auditory parabelt (areas CPB, RPB) | - | - | - | - | - | - | - |
| Superior temporal rostral cortex (area STR) | - | - | - | - | - | - | - |
| Superior temporal polysensory (areas TPO and PGa/IPa) | - | - | - | - | * | - | - |
| Inferior temporal (areas TE1-TE3 and TEO) | - | - | - | - | * | - | - |
| Visual extrastriate (areas V2, V4, V4T, 19M, MST, FSTd) | - | 0.11% | - | - | * | - | - |

\* indicates connections formed by fewer than 0.1% of the neurons participating in extrinsic connections.
- indicates no connection detected.
i indicates intrinsic connection (i.e. labeled cells located in the same area). These were not quantified due to the difficulty in distinguishing cells sending axons to the injection site versus those which incorporate tracer passively by diffusion through the extracellular medium.

**Table 3:** percentages of labelled neurons in different areas following injections in areas 24d (CJ167-CTBg) and 8b (other cases)

|  | **CJ167-**<br>**CTBg** | **CJ164-**<br>**FB** | **CJ83-**<br>**DY** | **CJ163-**<br>**DY** | **CJ164-**<br>**DY** |
| --- | --- | --- | --- | --- | --- |
| ***Granular frontal areas*** | **3.63%** | **15.87%** | **47.95%** | **45.41%** | **57.77%** |
| Frontal pole (area 10) | - | 1.23% | 2.20% | 0.69% | 6.20% |
| Dorsolateral prefrontal (areas 9, 46) | 1.07% | 1.37% | 13.04% | 8.69% | 31.82% |
| Ventrolateral prefrontal (areas 45, 47) | 0.20% | 3.60% | 2.40% | 4.08% | 3.93% |
| Orbitofrontal (areas 11, 13) | 0.32% | 5.66% | 0.77% | 1.79% | 4.96% |
| Area 8b | 0.45% | i<sup>a</sup> | i | i | i |
| Area 8aD | 1.42% | 2.44% | 28.76% | 27.25% | 10.66% |
| area 8aV | 0.16% | 1.56% | 0.78% | 2.91% | 0.20% |
| ***Medial prefrontal and anterior cingulate areas*** | **6.91%** | **18.74%** | **16.01%** | **19.88%** | **30.30%** |
| Medial prefrontal (areas 14, 32V, 32) | * | 2.66% | 10.43% | 8.90% | 26.61% |
| Ventral anterior cingulate (area 24a) | 0.16% | 1.38% | 2.30% | 1.98% | 0.43% |
| Dorsal anterior cingulate (area 24b) | 6.72% | 14.57% | 3.28% | 9.00% | 3.14% |
| Perigenual (area 25) | - | 0.12% | - | * | 0.12% |
| ***Motor and premotor areas*** | **49.56%** | **53.88%** | **25.69%** | **22.36%** | **3.79%** |
| Primary motor (area 4a-c) | 23.87% | 0.13% | * | * | * |
| Medial premotor (area 6M) | 11.15% | 13.99% | 7.57% | 0.63% | * |
| Dorsocaudal premotor (area 6DC) | 6.63% | 0.34% | 0.21% | 0.27% | * |
| Dorsorostral premotor (area 6DR) | 0.13% | 3.34% | 12.07% | 14.20% | 2.50% |
| Area 8C | 0.32% | 0.15% | - | 0.31% | * |
| Ventral premotor (areas 6Va, 6Vb) | 6.50% | 5.16% | - | 1.29% | - |
| Cingulate motor (areas 24c, 24d) | i | 17.89% | 5.74% | 4.69% | 0.14% |
| Proisocortical motor (area ProM) | 0.95% | 12.89% | * | 0.95% | 1.03% |
| ***Somatosensory areas*** | **10.93%** | **2.76%** | **0.01%** | **0.10%** | **0.03%** |
| Area 3a | 1.99% | 0.47% | - | * | - |
| Area 3b | 1.25% | 0.32% | - | * | - |
| Area 1-2 | 0.35% | 0.65% | - | * | - |
| Lateral somatosensory (Areas S2, S2PV, S2PR) | 3.20% | 1.21% | - | * | * |
| Somatosensory insular (areas GI, ReI) | 4.14% | 0.11% | * | * | * |
| ***Posterior parietal areas*** | **8.71%** | **0.23%** | **0.28%** | **0.44%** | **0.01%** |
| Rostral posterior parietal (area PE) | 1.96% | * | - | * | - |
| Medial posterior parietal (areas PGM, 31, 23c) | 3.17% | * | 0.13% | 0.13% | * |
| Visuomotor association (areas V6a, PEC, MIP, VIP, LIP, AIP) | 0.10% | * | * | * | * |
| Rostral ventral parietal (areas PFG, PF) | 2.61% | 0.11% | * | * | * |
| Caudal ventral parietal (areas OPt, PG) | 0.57% | * | * | 0.17% | * |
| Temporoparietal transition (area TPt) | 0.30% | - | * | * | - |
| ***Posterior cingulate and retrosplenial areas*** | **19.13%** | **2.56%** | **6.19%** | **9.62%** | **0.40%** |
| Area 29 | 0.58% | 1.11% | 0.71% | 1.29% | 0.28% |
| Area 30 | 2.77% | 0.93% | 3.24% | 2.91% | * |
| Area 23a | 4.49% | 0.44% | 1.64% | 4.31% | * |
| Area 23b | 11.29% | * | 0.37% | 0.92% | * |
| Areas 23V and Prostriata | - | - | 0.23% | 0.19% | - |

| ***Orbital and temporal limbic areas*** | ***1.03%*** | ***4.94%*** | ***0.37%*** | ***0.59%*** | ***5.16%*** |
| --- | --- | --- | --- | --- | --- |
| Gustatory cortex (area Gu) | 0.35% | 3.28% | - | 0.14% | 1.52% |
| Orbital and insular proisocortical areas (OPro, IPro, PaIM) | * | 0.46% | * | 0.26% | 2.59% |
| Orbital periallocortex (OPal) | * | 0.34% | * | 0.10% | 0.41% |
| Agranular and dysgranular insula | 0.55% | 0.18% | - | * | 0.02% |
| Piriform cortex | - | 0.15% | - | - | 0.27% |
| Entorhinal cortex | - | 0.17% | * | * | 0.13% |
| Perirhinal and temporal pole areas (areas 35, 36, TPro) | - | 0.34% | * | - | 0.20% |
| Parahippocampal (areas TH, TL, TF) | * | * | 0.17% | * | 0.02% |
| ***Temporal and occipital sensory areas*** | ***0.11%*** | ***1.03%*** | ***3.42%*** | ***1.59%*** | ***2.47%*** |
| Auditory lateral belt (areas CL, ML, AL) | - | - | 0.46% | - | - |
| Auditory parabelt (areas CPB, RPB) | - | - | 1.31% | * | * |
| Superior temporal rostral cortex (area STR) | - | * | 0.19% | - | 0.17% |
| Superior temporal polysensory (areas TPO and PGa/IPa) | * | 0.28% | 1.35% | 0.90% | 0.71% |
| Inferior temporal (areas TE1-TE3 and TEO) | * | 0.73% | * | 0.41% | 1.55% |
| Visual extrastriate (areas V2, V4, V4T, 19M, MST, FSTd) | * | * | * | 0.25% | * |

\* indicates connections formed by fewer than 0.1% of the neurons participating in extrinsic connections.
- indicates no connection detected.
i indicates intrinsic connection (i.e. labeled cells located in the same area). These were not quantified due to the difficulty in distinguishing cells sending axons to the injection site versus those which incorporate tracer passively by diffusion through the extracellular medium.

### Main observations following injections in cytoarchitectural area 6M

Figure 4 illustrates the combined pattern of cortical projections to area 6M, based on the registration of the neurons labelled by 5 injections of fluorescent tracers onto a volumetric representation of the Paxinos et al. (2012) stereotaxic atlas (Majka et al. 2016). Injections in sites attributed to area 6M revealed consistent characteristics in their pattern of afferents, as well as some variable features.

**Figure 4.**
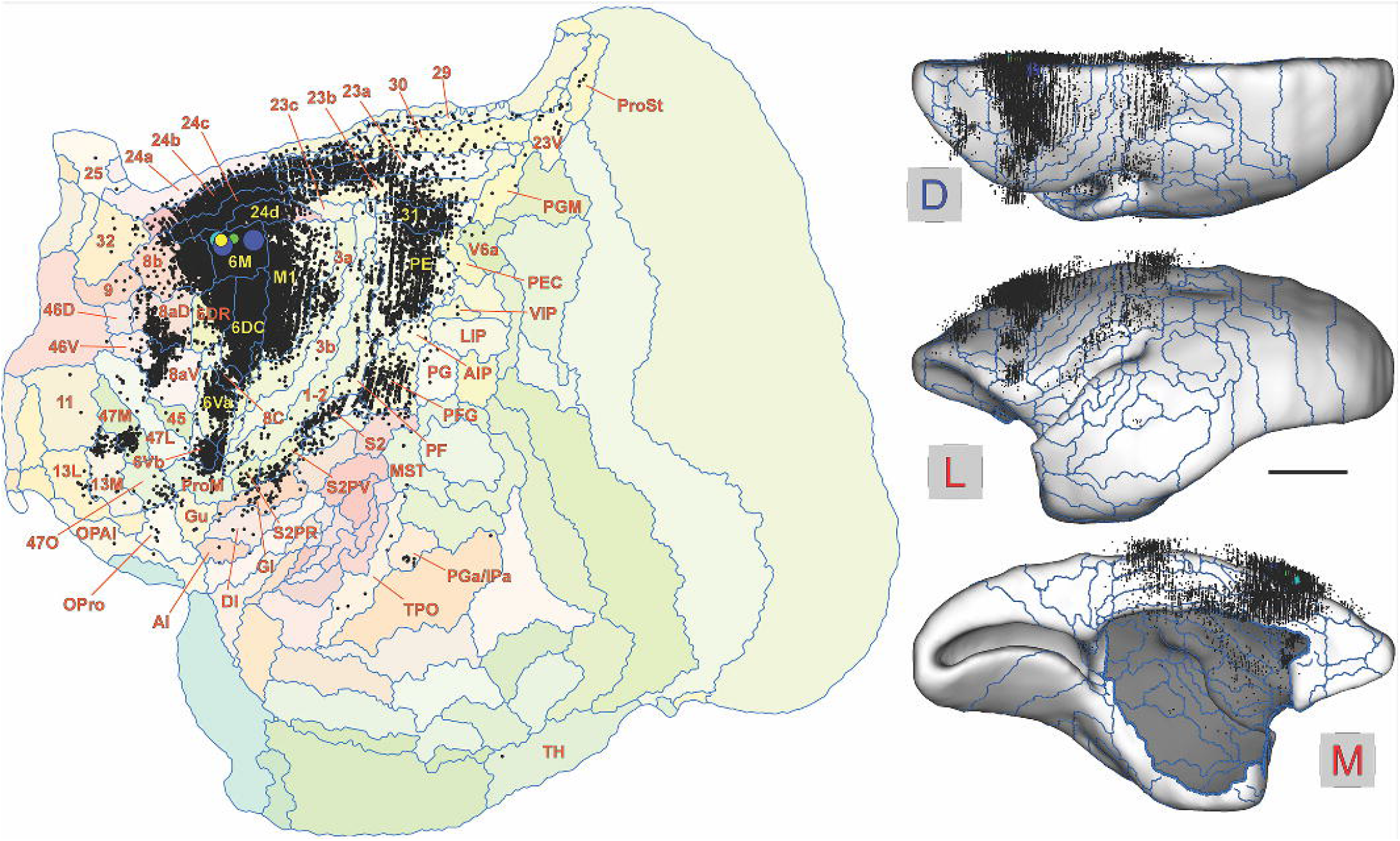
**Left:** Combined pattern of afferent projections to area 6M, based on computational co-registration of the results of 5 cases with fluorescent tracer injections to a flat map view of the Paxinos et al. (2012) brain template (Majka et al. 2016, 2020). In this representation, each black point represents a labeled neuron. Each of the areas containing at least 1 labeled neuron is identified according to the subdivision proposed by Paxinos et al. (2012; for abbreviations, see Supplementary Table 1). These maps provide convenient summaries of the distributions of labeled neurons in different areas, but, as with any such maps, there is some distortion due to different compression and stretch of different regions of cortex, which are inevitable due to intrinsic curvature (Chaplin et al. 2013). **Right:** dorsal (D), lateral (L) and medial (M) views of the brain (mid-thickness cortical surface) showing the locations of labeled neurons in stereotaxic space. Scale bar= 5 mm. Note that in the dorsal and lateral views only neurons located above the mid-thickness surface (i.e. towards the supragranular layers) become visible in this representation. In the medial view, some infragranular neurons are visible in the region where the corpus callosum and subcortical structures were removed.

In all cases, the majority (66.6 ± 8.8%, mean and s.d.) of afferents originated from other motor and premotor areas, in particular the primary motor cortex (M1, cytoarchitectural fields 4a-c of Paxinos et al. 2012; 14.6 ± 12.9%), the dorsocaudal (6DC) and dorsorostral (6DR) premotor areas (18.7 ± 6.6% and 8.5 ± 5.9%, respectively), and the putative cingulate motor areas (24c and 24d of Paxinos et al. 2012; 21.7 ± 3.7%). Fewer and less consistent projections originated in area 8 caudal (8C, a strip of putative oculomotor cortex which separates dorsal and ventral premotor areas; Burman et al. 2015), the ventral premotor areas (primarily, cytoarchitectural area 6Va) and the proisocortical motor area (ProM), which has been linked to the motor control of vocalizations (Alipour et al. 2002).

Neurons forming connections with area 6M were also present in several prefrontal areas. The most consistent among these projections originated in subdivisions of area 8 complex (Burman et al. 2006; Reser et al. 2013), including in particular area 8b (10.0 ± 11.2%) and the dorsal part of area 8a (8aD; 1.0 ± 1.6%). Other prefrontal connections were observed mostly following injections in the rostral part of 6M (see below). Consistent long-range connections were also observed with anterior cingulate area 24b (4.4 ± 2.9%), posterior parietal area PE (area 5; 5.7 ± 6.1%) and posterior cingulate areas 23a and 23b (1.3 ± 0.7% and 1.1 ± 0.6%, respectively). The mesial posterior parietal cortex also contributed significant contingents of long-range projections (2.5 ± 1.7%), of which the majority originated in area 31 and area 23c (which includes the putative marmoset homologue of the macaque’s transitional somatosensory area, TSA; Morecraft et al. 2004).

Although the spatial distribution of projection neurons was similar in all cases with injections in 6M, we also observed variations in the relative proportions of projections according to rostrocaudal location (Fig. 5). As detailed below, caudal injections (e.g. cases 1-3) were characterized by a paucity of prefrontal afferents, and stronger projections from M1, somatosensory areas, and posterior parietal areas, in comparison with rostral sites (e.g. cases 5-7). Case 4 (CJ-113FE) was, in many ways, intermediate, showing characteristics observed in both caudal and rostral injections (Table 2). These trends are also illustrated in Suppementary Figure S1.

**Figure 5.**
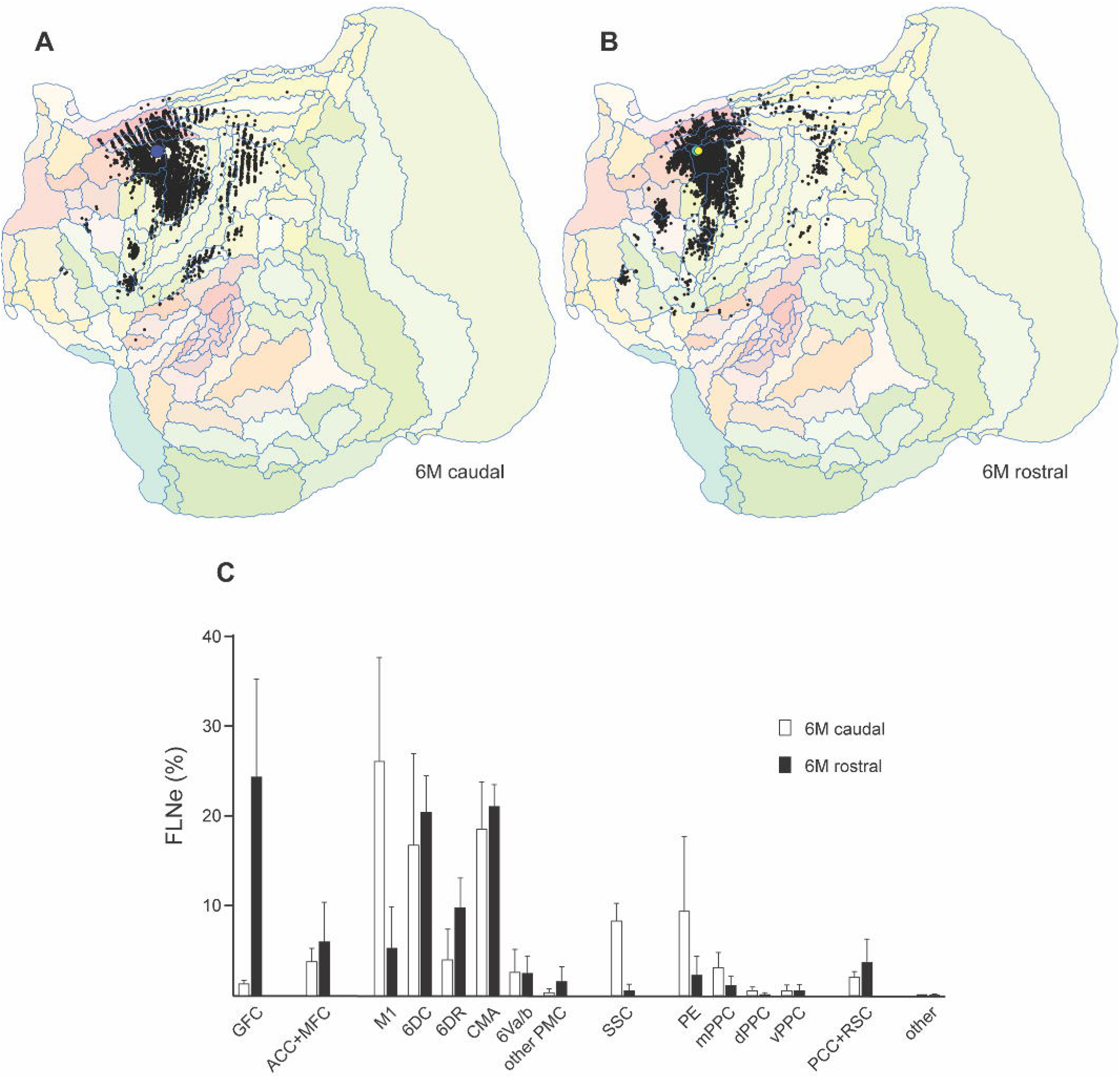
**A:** distribution of labeled neurons following an injection in caudal area 6M (case 1, CJ94-FB), shown in a flat map of the marmoset cortex. **B:** combined distribution of labeled neurons following 2 injections in rostral area 6M (cases 6 and 7, CJ170-DY and CJ164-CTBg). Data from cases 2 and 3 were not included due to the lack of availability of computational registration to the template, case 4 was not included because it showed transitional characteristics between caudal and rostral injections, and case 5 was not included given the large number of labeled neurons (Table 1), which would make visual comparison with caudal 6M injections difficult. Thus, this comparison involves similar numbers of labeled neurons outside area 6M (4830 vs. 4551 cells; Table 1). This comparison shows that the spatial distribution of neurons was similar between injections at the rostral and caudal ends of area 6M, despite quantitative differences. **C:** Comparison of the percentages of labeled neurons in different cortical areas and regions. The white bars represent the mean and standard deviation of the FLNe (fraction of labeled neurons, extrinsic connections) in cases 1-3 (caudal 6M injections), and the black bars represent the same quantities for cases 5-7 (rostral 6M injections). Abbreviations represent areas or groupings of areas according to Table 2: GFC – granular frontal cortex (areas 8, 9, 10, 11, 13, 45, 46, 47 and their subdivisions); ACC+MFC –anterior cingulate cortex and medial frontal cortex (areas 14, 24a, 24b, 25, 32 and 32V); M1 – primary motor cortex (areas 4a, b, c); 6DC – area 6, dorsocaudal; 6DR – area 6, dorsorostral; CMA-cingulate motor areas (areas 24c, 24d); 6Va/b – ventral premotor areas; other PMC – other premotor cortex (areas 8C [8 caudal] and ProM [proisocortical motor]); SSC – somatosensory cortex (areas 1-2, 3a, 3b, subdivisions of the second somatosensory complex [S2], and insular somatosensory (GI [granular insular] and ReI [retroinsular]); PE – parietal area PE; mPPC – medial posterior parietal cortex (areas 23c, 31 and PGM); dPPC – dorsal posterior parietal cortex (areas AIP, LIP, MIP, VIP, PEC and V6a); vPPC – ventral posterior parietal cortex (areas PF, PFG, PG, OPt and TPt); PCC+RSC – posterior cingulate cortex and retrosplenial cortex (areas 23a, 23b, 23V, 29, 30 and ProSt [prostriata]).

### Projections to caudal 6M (cases 1-3)

The patterns of connections revealed by an injection in the caudal part of area 6M (case 1, CJ94-FB) is illustrated in detail in Figures 5A and 6. Although the percentages of labeled neurons varied (Table 2), the pattern of projections revealed in this case was very similar to that observed in cases 2 and 3. Most cortical projections to this injection site (80% of the extrinsic afferents) originated in other areas implicated in motor control, including in particular M1 (Fig. 6C, D), the putative cingulate motor areas (24c/d; Fig. 6A-D) and area 6DC (Fig. 6B, C). As for cases 2 and 3 (Table 2), afferents from area 6DR were much less numerous (Fig. 6A, B). Ventral premotor afferents originated primarily from the dorsocaudal area (6Va), but also included the ventrorostral area (6Vb; Fig. 6B). Afferents from the proisocortical motor area (ProM) and area 8C were very sparse and inconsistent among rostral 6M injection cases (Table 2).

**Figure 6.**
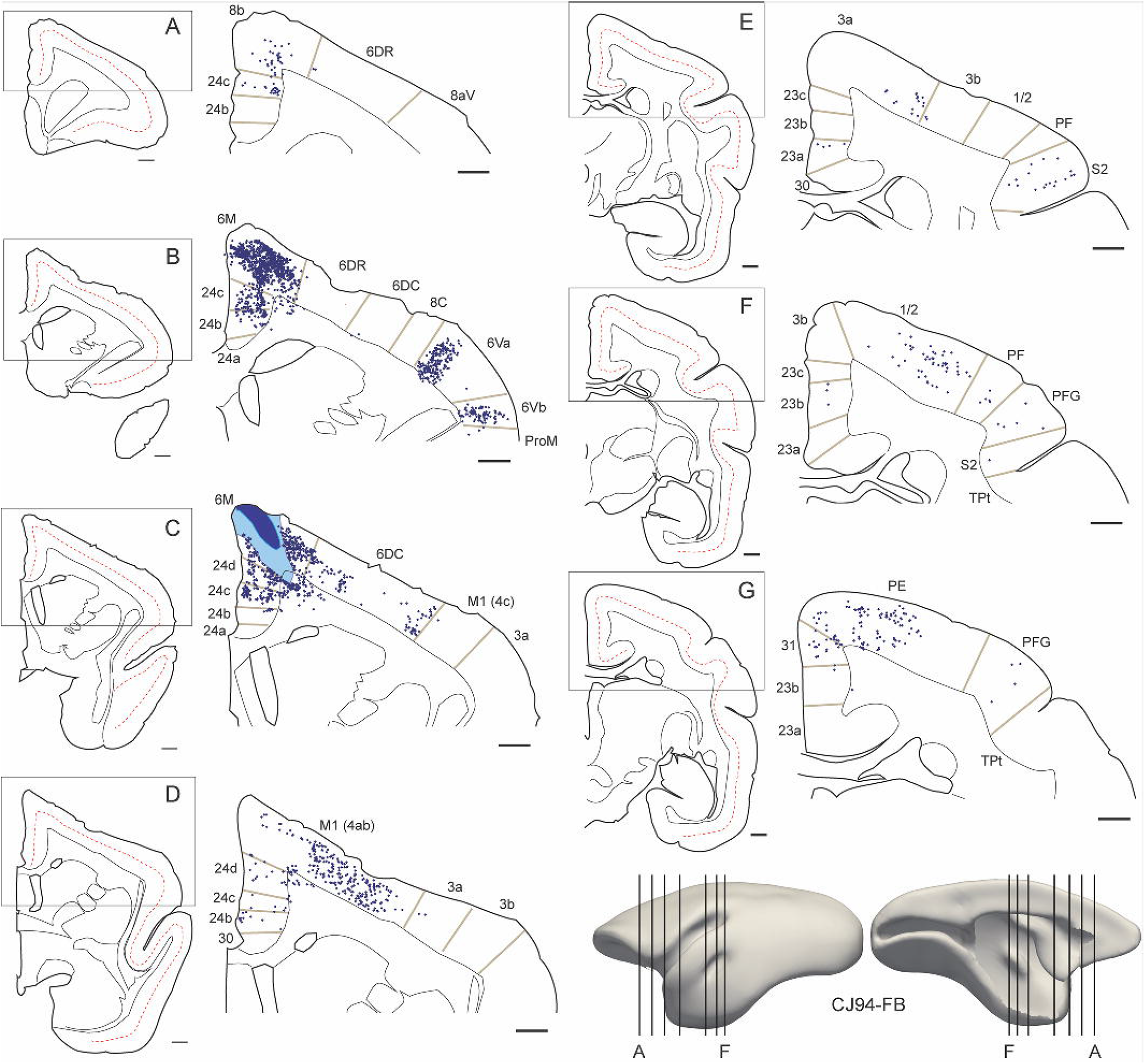
**A-G:** Representative coronal sections showing the locations of labeled cells after injections in the caudal part of 6M in case 1 (CJ94-FB). The levels of the sections are indicated in 3-D views of the population-average marmoset left hemisphere (bottom right; from Majka et al. (2021). Each panel illustrates a low power view of the section (left) and a magnified view of the boxed region, containing labeled neurons (right). On the left panels, the center of layer 4 is indicated by the dashed red line. The right panels illustrate the locations of single labeled neurons (dark blue circles) relative to cyto- and myeloarchitectural borders (grey lines). Only areas containing labeled neurons and adjacent areas are identified (for abbreviations, see Supplementary Table 1). In C, the FB injection site is illustrated (dark blue – core; light blue – halo). A scale bar (1 mm) is provided on the bottom right of each section drawing.

Prefrontal projections to caudal 6M originated primarily in area 8b (Fig. 6A). Another major source of afferents in the frontal lobe was cingulate area 24b (Fig. 6B-D), with cingulate area 24a also contributing sparser connections.

In the parietal cortex, somatosensory areas formed an important group of projections to caudal 6M, with afferent neurons located in rostral parietal areas 3a and 1-2 (Figs 5A and 6D-F), but less densely in the primary somatosensory area (3b). Labeled cells were also observed in the shoulder and dorsal part of the lateral sulcus (S2 complex; Fig. 6E-F) and, in small numbers, in the granular insular cortex. Posterior parietal afferents were concentrated primarily in the rostral part of area PE (Fig. 6G), ventral parietal areas PF and PFG (Fig. 6F, G), and medial areas 31 (Fig. 6G) and 23c. Finally, posterior cingulate afferents were observed in the rostral parts of areas 23a and 23b (Fig. 6E-G), but very few neurons were found in areas 29 and 30 (see also Table 2).

### Projections to rostral 6M (cases 5-7)

The detailed pattern of connections in one rostral 6M injection case is shown in Figure 7 (case 170-DY). The spatial distribution of labeled neurons was similar to that observed after injections in the caudal part of area 6M (Fig. 5A, B), but subtle differences emerged from quantitative analysis (Fig. 5C).

**Figure 7.**
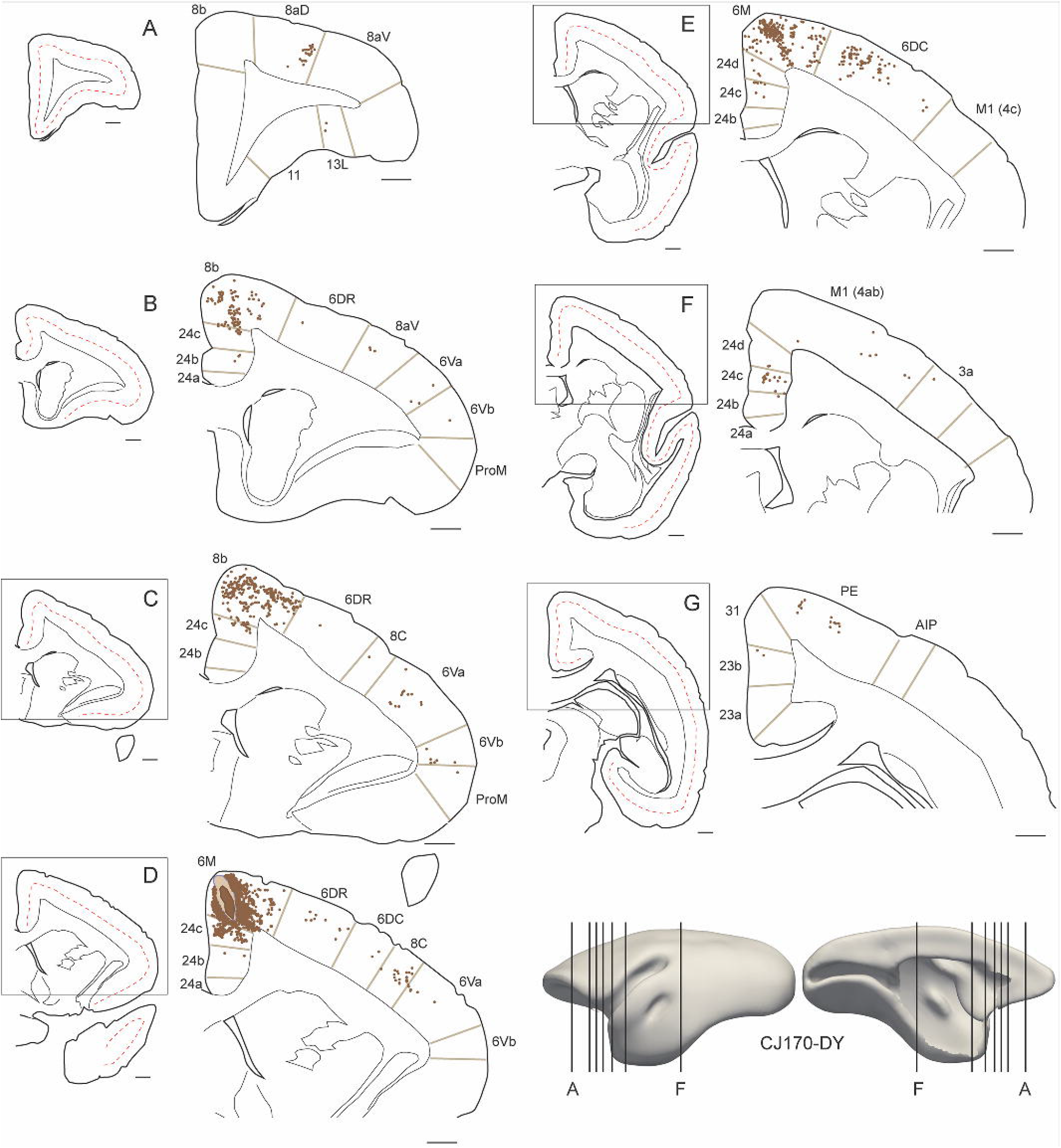
**A-G:** Representative coronal sections showing the locations of labeled cells after injections in the rostral part of 6M in case 6 (CJ170-DY). The levels of the sections are indicated in 3-D views of the population-average marmoset left hemisphere (bottom right; from Majka et al. (2021). Each panel illustrates a low power view the section (left) and a magnified view of the boxed region, containing labeled neurons (right). On the left panels, the center of layer 4 is indicated by the dashed red line. The right panels illustrate the locations of single labeled neurons (brown circles) relative to cyto- and myeloarchitectural borders (grey lines). Only areas containing labeled neurons and adjacent areas are identified (for abbreviations, see Supplementary Table 1). In **D**, the DY injection site is illustrated (dark brown – core; light brown – halo). A scale bar (1 mm) is provided on the bottom right of each section drawing.

Caudal frontal lobe areas involved in motor planning and control also supplied the bulk of extrinsic projections (mean ± s.d.: rostral 6M, 60.6 ± 2.1% vs. caudal 6M, 70.7 ± 11.3%). However, compared to the cases with caudal injections, there was a decreased emphasis in projections from M1 (Fig. 7E, F). Unlike in caudal 6M injections (e.g. Fig. 5A), the vast majority of M1 afferent cells was located in the rostral half of this area (Fig. 5B). Conversely, rostral 6M received relatively more afferents from area 6DR (Figs. 5C and 7C, D, Table 2).

A second significant difference was the percentage of projections from the granular frontal cortex (rostral 6M, 24.4 ± 10.8% vs. caudal 6M, 1.3 ± 0.5%; see Table 2 and Figure 5C). Despite the quantitative difference, the locations of the main groups of afferent neurons were similar, with clusters of cells concentrating in the caudal half of cytoarchitectural area 8b (Fig. 7B, C), the rostral portion of the border between dorsolateral prefrontal areas 8aD and 8aV, and the rostral part of orbitofrontal area 13L (Figs. 5A, B and 7A). Small numbers of labeled neurons were found in area 9 and subdivisions of area 47, primarily in case 5 (CJ163-FB), which was relatively large and reached farthest into layer 6 (Fig. 1), leading to the largest number of labeled neurons overall.

In comparison with caudal injections, rostral 6M injections also revealed reduced numbers of neurons in somatosensory (8.2 ± 2.1% vs. 0.6 ± 0.7%) and posterior parietal (13.8 ± 9.3% vs. 4.5 ± 3.3%) areas. For example, case 1 (Fig. 5A) shows numerous clusters of labeled cells across areas 3a, 3b, 1-2 and the S2 complex, whereas the combined pattern from the 2 injections shown in figure 5B only shows scattered cells in the facial and oral representations in areas 3a and 1-2 (e.g Fig. 7F). The main cytoarchitectural areas in the posterior parietal cortex providing the afferents are the same in rostral and caudal cases (namely, lateral parietal areas PE, PFG, and PF, and medial parietal areas 31 and 23c), although in PE (Fig. 7G) the cells forming the connections were located on average more caudally (compare Fig. 5A, B); in addition, these projections were very sparse following the most rostral injection (CJ164-CTBg, case 7). Finally, as for caudal 6M, projections from the posterior cingulate cortex originated in areas 23a, 23b and 30 (Figs. 5B, 7G).

### Comparison with area 8b

The nature of the transition from area 6M to the dorsomedial prefrontal cortex (area 8b; Reser et al. 2013) has not been explored in detail. Therefore, a natural question is whether the connectional differences we observe between rostral and caudal 6M are compatible with an alternative model whereby the sites we attributed to rostral 6M are actually best seen as part of area 8b. The present dataset, including 4 injections in the territory assigned to 8b, provided an opportunity to explore this question. As hinted by Reser et al. (2013), the sources of cortical afferents to area 8b were somewhat variable in terms of the specific areas, an observation that could suggest either internal subdivisions of this area, or a type of modular organization in connectivity. Nonetheless, as detailed below, some common observations distinguish area 8b from area 6M. Figure 8 illustrates the combined pattern of labeled neurons following the injections in area 8b, and Figure 9 (A-D) illustrates individual projection patterns (cases 8-11) and a summary view of the distributions of labeled extrinsic neurons following injections in this area, using the same categories used in Figure 5. In addition, Table 3 provides quantitative information about the percentages of labeled neurons in different areas.

**Figure 8.**
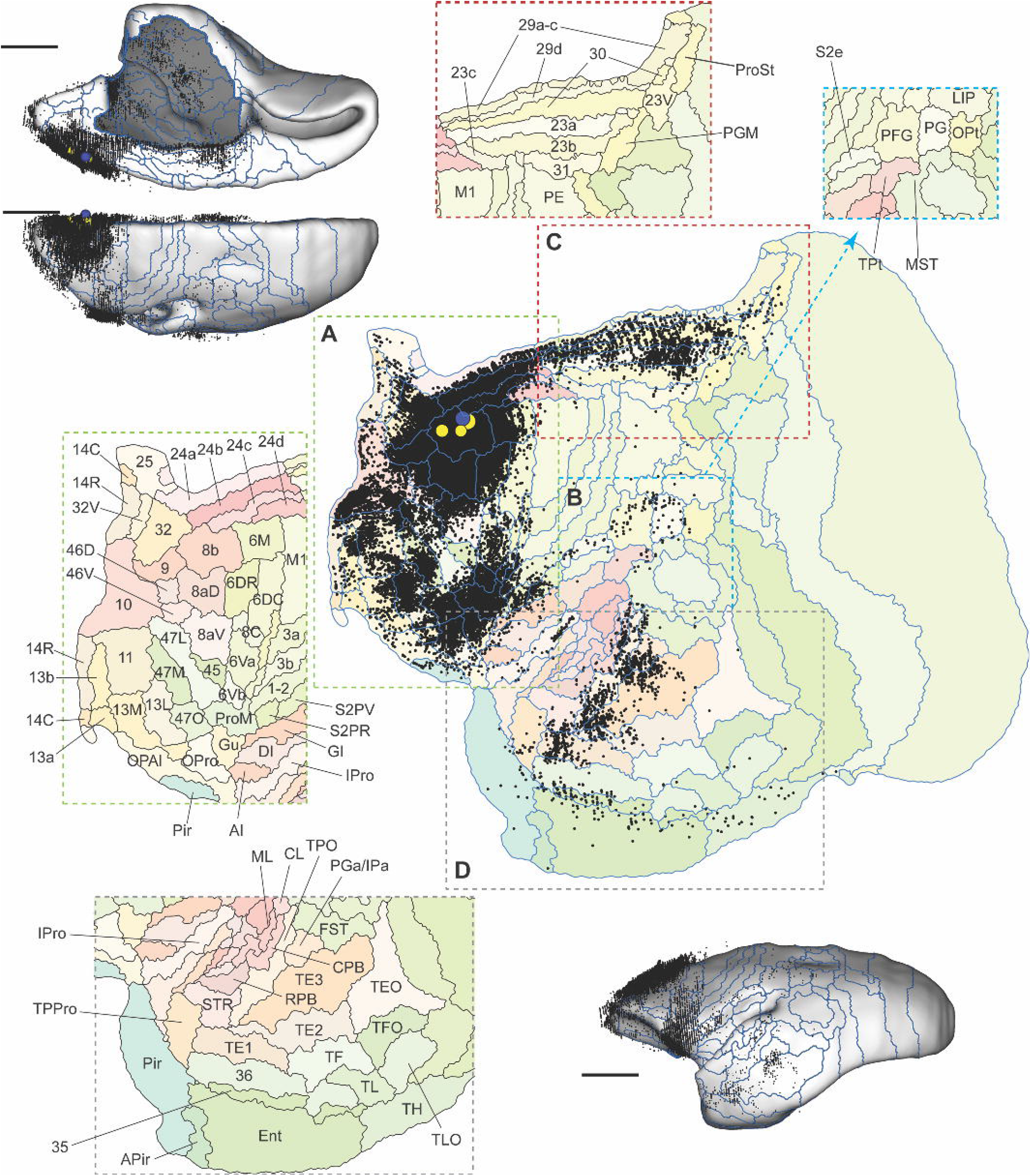
Combined pattern of cortical projections to area 8b, based on the co-registration of four tracer injections (cases 8-11). The main panel is a flat map representation of the marmoset left cortex (based on Paxinos et al. 2012), showing the location of the injections (blue and yellow circles) and labeled neurons (each black point represents a single cell). Comparison with the pattern shown in Figure 4 demonstrates the differences relative to area 6M. Given the density of labeled neurons, the areas containing at least 1 labeled cell are identified in adjacent boxes representing the frontal lobe (**A**), lateral parietal cortex (**B**), posterior cingulate and retrosplenial cortex (**C**) and temporal lobe (**D**). The inserts illustrate medial, dorsal (top left) and lateral (bottom right) views of the brain (mid-thickness cortical surface) showing the locations of labeled neurons in stereotaxic space. Scale bars= 5 mm. Note that in the dorsal and lateral views only neurons located above the mid-thickness surface (i.e. towards the supragranular layers) become visible in this representation. In the medial view, some infragranular neurons are visible in the region where the corpus callosum and subcortical structures were removed.

**Figure 9.**
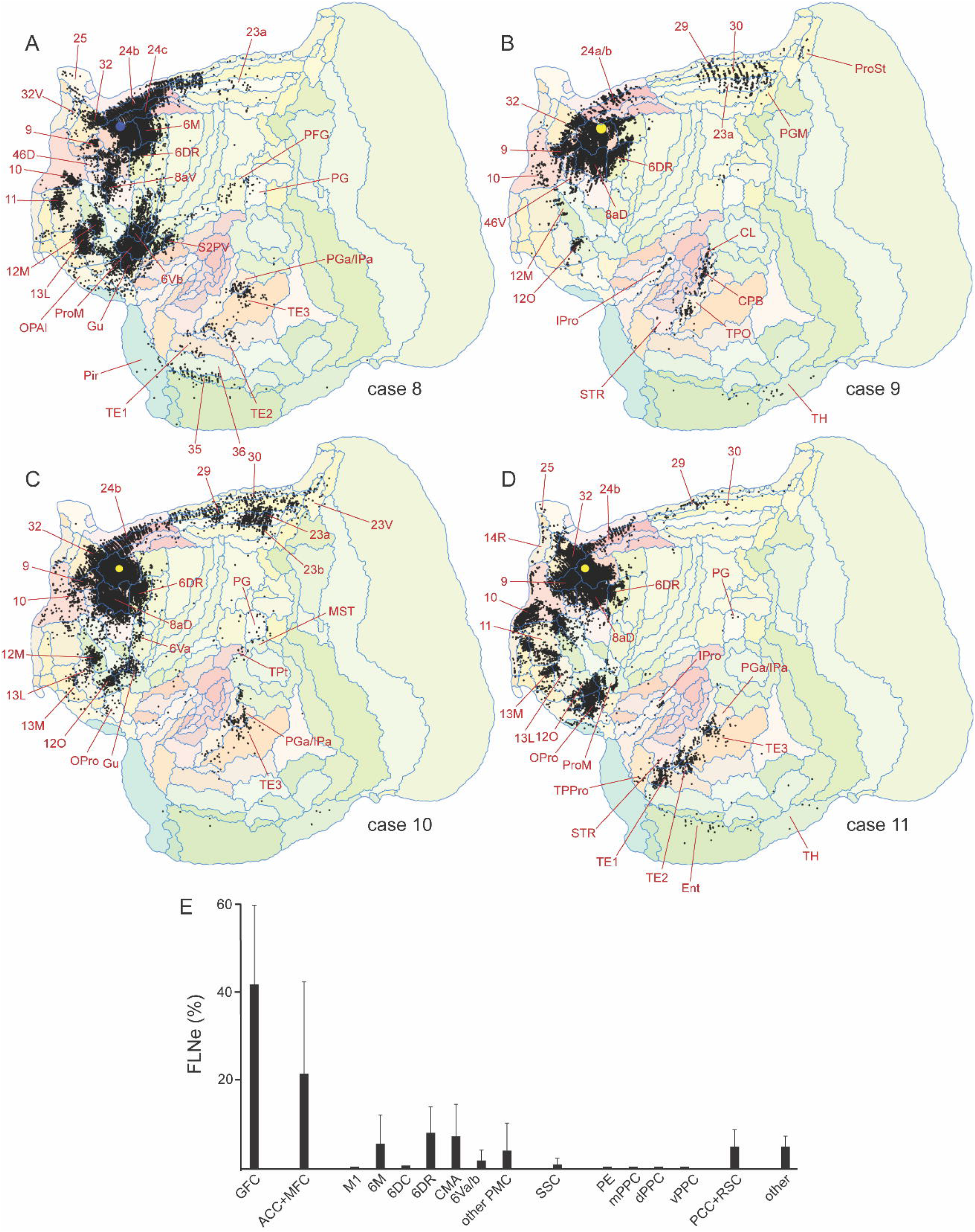
**A-D:** Individual flat maps showing the distribution of labeled neurons in 4 cases with injections in area 8b. See also legend to Figure 8. **E:** Percentages of labeled neurons (extrinsic connections) for area 8b injections, grouped in the same categories used in Figures 5 and 10.

These analyses reveal that the sources of afferents to area 8b are quite different from those found following injections in rostral 6M. For example, there are relatively few afferents from areas involved in motor planning or control (mean, s.d: 21.2 ± 6.3%; Table 3), with most labeled cells being located in areas 6M, 6DR, 24c and 6Vb (Fig. 8A). Conversely, prefrontal afferents were widespread (Fig. 8A), including for example dense projections from areas 9 and 10 (see Burman et al. 2011 for reciprocal connections). Projections to area 8b also included most of the dorsolateral, ventrolateral, orbitofrontal and rostromedial prefrontal areas (Burman et al. 2006; Burman and Rosa 2009), many of which were weakly or not labeled following area 6M injections (e.g. areas 11, 12, 13, 14, 32, 46 and its subdivisions). In the anterior cingulate cortex, the label was concentrated in area 24b, but sparse in area 24a (Fig. 8A).

Label in the posterior parietal cortex (Fig. 8B) was much sparser (0.24 ± 0.18%; Table 3) in comparison with that observed after injections in area 6M (9.14 ± 7.35%; Table 2), with the emphasis shifted caudally, towards areas PG (Fig. 9 A, C, D), TPt and PGM (Fig. 9B, C). Notably, area PE was nearly devoid of label, in sharp contrast with injections in area 6M. In the posterior cingulate and retrosplenial regions (Fig. 8C) label was also shifted caudally relative to projections to area 6M, not only with respect to the extents of areas 23a and 23b, but also by including visual association areas 23V and prostriata (e.g. Fig. 9B, C; Palmer and Rosa 2006b; Yu et al. 2012), and by showing denser projections from subdivisions of area 29. In addition, multiple areas in the caudal orbital, insular and ventral temporal limbic cortex formed projections to area 8b (Figure 8A; Fig. 9), in different proportions depending on the location of the injection sites (Table 3, and Fig. 9B-D). Overall, the contribution of projections from these areas was more substantial (mean, s.d: 2.76 ± 2.64%) than that revealed by injections in area 6M (Table 2). Notably, 3 of the injections revealed connections from the putative gustatory cortex (Burman and Rosa 2009; see Table 3 and Fig. 9A, C, D).

Finally, unlike injections in area 6M, those in area 8b consistently labeled neurons across multiple temporal lobe areas involved in higher-order sensory processing (Fig. 8D). Although projections from any individual area contributed relatively small percentages of the extrinsic afferents to area 8b (Table 3), they originated from a wide variety of areas, with emphasis on the superior temporal polysensory cortex (areas TPO and PGa/IPa) and the dorsalmost part of cytoarchitectural area TE, which in macaques has also been reported to harbor neurons with polysensory properties (Baylis et al. 1987). The label following some of the injections (Fig. 9A, C, D) included the dorsal subdivision of the medial superior temporal visual area (MST) and the fundus of superior temporal area (FST), where motion-selective visual responses are observed (Rosa and Elston 1998). In addition, several injections also revealed afferents from auditory association areas, emphasizing the parabelt and superior temporal rostral cortex. Projections from the entorhinal, ectorhinal (35), perirhinal (36) and piriform (Fig. 9A, D) areas were also observed. These results hint at a role of area 8b in multisensory integration, where closely located sites receive different combinations of sensory input.

### Comparison with adjacent motor and premotor areas

We were also interested in gaining insight into how the subdivisions of area 6M differ from adjacent areas involved in motor control and planning. For this purpose, we compared the connection patterns of 6M with those previously reported for the marmoset areas M1, 6DC and 6DR (Burman et al. 2014a, b) and with new observations regarding the connections of the dorsal part of the cingulate motor cortex (area 24d; case 12, CJ167-CTBg). Figure 10A illustrates the combined pattern of connections revealed by 5 injections in the dorsal part of area M1, where representations of the axial musculature and limbs were observed (Burman et al. 2008). Figures 10B and 10D show similar representations for the connections of areas 6DC and 6DR (4 injections each), and the pattern observed after a single injection in area 24d is shown in Fig. 10C.

**Figure 10.**
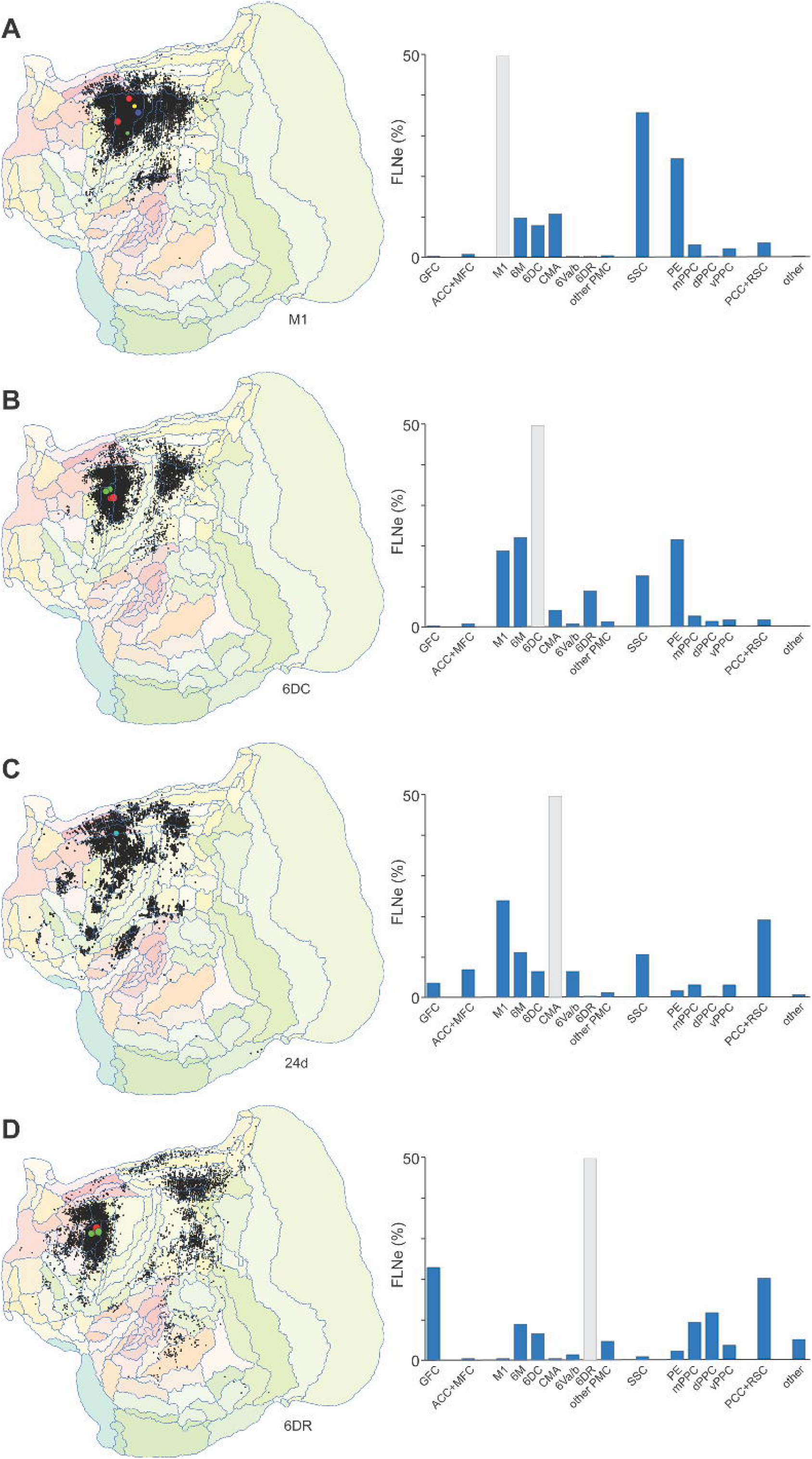
Comparison of the patterns of label resulting from injections in areas adjacent to area 6M. Each panel (**A-D**) shows the location of labeled neurons (left) and the fraction of extrinsic labeled neurons (FLNe, right). In the right panels, grey bars indicate the area containing the injections, in a format similar to that used in Figure 5 to illustrate the connections of area 6M (see legend of Figure 5 for abbreviations). Data shown in panels A, B and D are from Burman et al. (2014a, b, 2015), and panel C represents case 12 of the present study **A:** composite pattern revealed by 5 injections in the medial part of the primary motor area (M1). **B:** 4 injections in the medial part of the dorsocaudal premotor area (6DC). **C:** 1 injection in area 24d (part of the cingulate motor complex); **D:** 4 injections in the medial part of the dorsorostral premotor area (6DR).

This analysis highlights the uniqueness of M1 in the motor control network, particularly with respect to the robust connectivity with the somatosensory cortex (SSC, Fig. 10A, right).; on average, as many as 35.8% of the afferents stemming from the rostral (areas 3a, 3b, and 1-2, the traditional “S1”) and lateral (in particular, area S2) parietal complexes. This is evident from the combined flat map (Fig. 10A, left) in that the projecting neurons occupied a continuous region linking the premotor and posterior parietal areas. In contrast, for all other areas explored (Figs. 4, 5 and 10B-D), there is a noticeable gap in label in the region corresponding to the traditional S1. M1 also stood out by the numerical strength of the afferents from posterior parietal area PE (mean 24.6% of extrinsic connections) and by the fact that the vast majority of other connections originated from other premotor areas, in particular 6M, 6DC and the cingulate motor areas (24c/d), which contributed approximately equal numbers of projecting neurons.

The second tier of motor-related areas (i.e. areas that share a border directly with M1) included the caudal subdivision of area 6M (Figure 5A, C) and areas 6DC and 24d (Fig. 10B, C). These areas share the characteristic of receiving major inputs from M1 (on average, ∼20-30% of their extrinsic connections) and from other premotor areas, as well as relatively robust afferents from somatosensory areas (8-13%). At the same time, each of these areas shows specific emphases in their connectivity. For example, both 6DC and caudal 6M show relatively strong connections with parietal area PE (21.7% and 9.5% of the afferents, respectively) and rostral premotor area 6DR (9.1% and 4.0%), in comparison with area 24d (PE: 2.0%; 6DR: 0). Conversely, area 24d stands out relative to 6DC and caudal 6M by virtue of the numerous afferents it receives from posterior cingulate areas 23a, 23b and 30 (19.3%) and anterior cingulate area 24b (6.9%). Other notable differences include the presence of major inputs from the granular insular cortex to 24d, which correspond to a relatively large fraction of the somatosensory-related afferents to this area (4.1%), but not to 6DC or caudal 6M (Table 3).

Finally, the rostral part of area 6M and area 6DR shared key similarities which hinted at a different level of involvement in premotor function, relative to caudal 6M, 6DC and 24d. Most notably, these areas share a relatively more robust input from granular frontal areas (GFC in Figs. 5C and 10D; rostral 6M: 24.4%; 6DR: 23.3%), which originated in largely overlapping regions (Figs. 5B and 10D), and very sparse somatosensory afferents. They both receive significant inputs from other motor and premotor areas, albeit in different proportions: for example, the direct input from M1 to rostral 6M is far more significant than that to 6DR (5.2% vs. 0.1%), and rostral 6M is strongly connected to the cingulate motor areas (24c/d). In the parietal lobe, the projections to 6DR originate from a wide diversity of areas in the medial, dorsal and ventral regions, whereas the input to rostral 6M is more restricted, being primarily from areas PE, 31 and PF/PFG. Finally, unlike rostral 6M, area 6DR receives significant afferents from superior temporal polysensory areas and areas 29 and 30.

## Discussion

We used the pattern of cortical connections to investigate the extent to which marmoset area 6M, proposed largely based on location and cytoarchitecture (Burman et al. 2006; Burman et al. 2008; Paxinos et al. 2012), is anatomically similar to the medial premotor areas described in other primates. We were particularly interested in establishing if 6M encompassed subregions along the rostrocaudal dimension, which could be homologous to the SMA and preSMA (F3 and F6; Matelli et al. 1991). Finally, we explored the connectivity differences that distinguish area 6M from its neighboring subdivisions of prefrontal and motor-related cortex. The present results complement earlier findings (Burman et al. 2014a, 2014b; 2015; Bakola et al. 2015) in providing a quantitative, network view of the motor-related cortex in the marmoset.

### Comparative analyses

We found that area 6M receives numerous projections from the primary motor and premotor areas, including the putative cingulate motor cortex, as well as highly specific projections from anterior cingulate, prefrontal, parietal somatosensory and association areas, and posterior medial areas. Our main results are in agreement with earlier studies on the connectivity of the medial part of cytoarchitectural area 6 in macaques (McGuire et al. 1991; Luppino et al. 1993; Morecraft and Van Hoesen 1993; Morecraft et al. 2012), squirrel monkeys (Jürgens1984) and prosimians (Fang et al. 2005), and are also partially consistent with data from a human post-mortem analysis (Vergani et al. 2014), with regards to motor and midline connections.

Previous anatomical tracing studies in marmosets have reported afferents originating from the 6M region (Krubitzer and Kaas 1990; Huffman and Krubitzer 2001; Roberts et al. 2007; Reser et al. 2013; Burman et al. 2014a, 2014b; Burman et al. 2015), and our results showed reciprocal pathways to most, if not all of these. In addition, although tracer injections have not directly targeted the medial premotor cortex in other New World monkeys, the spatial distribution of connections revealed by injections in other areas also suggest a similar spatial distribution (Stepniewska et al. 1993; Padberg et al. 2005; Gharbawie et al. 2011). Thus, one of our main conclusions is that the organization of the medial premotor cortex in marmosets is essentially similar to that of other primates, despite variations in the motor repertoire among species. This, in turn, indicates marmosets can be valuable animal models for studying the mechanisms involved in motor planning and execution, particularly given the opportunities represented by the lissencephalic configuration of the frontal cortex. Finally, even though all 6M sites showed a relatively consistent projection pattern, we found a gradient in the strength of inputs (with sensorimotor and prefrontal afferents preferentially targeting caudal and rostral parts of 6M, respectively), which we interpret as providing evidence that marmoset area 6M is likely to include both the SMA and preSMA. It is important to acknowledge, however, that the latter results do not exactly replicate thoser of earlier work in macaques, where the preSMA is reported not to receive afferents from M1 (Luppino et al. 1993). However, as discussed below, detailed comparison of the present observations with those of previous studies requires careful prior reflection on the likely effect of the different brain (and cortical area) volumes in the results of anatomical tracing experiments. At least two factors need consideration in this respect.

First, given that the macaque brain is at least 12 times larger (in volume) than the marmoset brain, similar-sized tracer injections will necessarily occupy a much larger fraction of any given cortical area in the marmoset, compared to the macaque. Thus, the likelihood that an injection is entirely restricted to the pre-SMA, which is small relative to the SMA, is necessarily lower in the marmoset. Given this, we consider the present results as inconclusive with respect to differentiating between a model whereby cytoarchitectural area 6M is in fact comprised of two distinct fields with a defined border (Luppino et al. 1993), and another in which this area comprises a functional gradient (where the proportions of modules receiving different combinations of afferents vary; see Rosa and Tweedale 2005; Graziano and Aflalo 2007 for discussion). In general, these two possibilities cannot be distinguished without an experimental design that includes injections that are much smaller than the area(s) being explored, and a comprehensive grid of injections. Connectional gradients have been observed within visual areas with sharp architectural borders (Palmer and Rosa 2006b; Majka et al. 2019), and have also been proposed for parietal association areas (e.g. MIP and PGM; Bakola et al. 2017, Passarelli et al. 2018).

Second, theoretical considerations lead to the expectation that larger brains will show decreased “connectedness” in comparison with smaller brains (Ringo 1991; Striedter 2005), due to factors such as limited capacity for the synaptic space in the neuron’s membranes to increase in proportion to the number of neurons. Evidence in support of the validity of this principle has been found in the connections of the frontal pole cortex (Rosa et al. 2019) and visual area MT (Palmer and Rosa 2006a). In both cases, homologous areas were found to share the same principal connections in different primate species, but to also receive additional sparse connections in the smallest brain (marmoset). Most of these additional connections exist as dysynaptic pathways in larger brains, such as those of macaques. Thus, the likelihood of allometric scaling of connections also needs to be taken into consideration when interpreting the significance of discrepancies such as the existence of projections from M1 to rostral 6M in the marmoset, in comparison with the reported absence (Luppino et al. 1993) or paucity (Morecraft et al. 2012) of such connections to the macaque pre-SMA (F6).

In summary, while the present results are *compatible* with a fundamental similarity in organization of the medial premotor cortex across primates, they also highlight the fact that species differences are not only possible, but likely. Further, the possibility should be considered that the SMA and pre-SMA in the marmoset are less differentiated from each other than in the macaque. In the latter species a cytoarchitectural difference between these areas has been described, linked to the fact that the former contains larger pyramidal cells in layer 5, which form corticospinal projections (Luppino e al. 1994). Although this difference was not obvious in the present analysis (which is based on the criteria of Paxinos et al. 2012), a more focused study where cytoarchitecture is correlated with the results of retrograde tracers in the spinal cord and additional staining techniques may reveal subtle differences.

### Topographic organization of 6M

The lack of microstimulation maps remains a limitation to the precise comparative interpretation of the functional organization of the medial premotor cortex in the marmoset. To date, the definition of premotor areas in this species has relied on cytoarchitectural and myeloarchitectural features (Burman et al. 2006; Paxinos et al. 2012), with data on connections providing support for the proposed homologies (Burman et al. 2014b). Although two studies (Burish et al. 2008; Burman et al. 2008) demonstrated that microstimulation of the caudal part of area 6M can elicit movements, these results were not sufficient to derive a comprehensive view of the topographic organization of this cytoarchitectural area.

In the macaque SMA movements of the hindlimb, forelimb and facial musculature are found in caudal to rostral sequence, with the last occupying a relatively small sector (Luppino et al. 1993). In addition, stimulation of the pre-SMA mostly results in arm movements. A similar somatotopy within the marmoset cytoarchitectural area 6M is suggested from the topographic organization of its connections with M1 (Burman et al. 2014a). Retrograde tracer injections in the representation of the hindlimb and axial musculature of M1 revealed projections from caudal 6M, including the border with M1 (their cases 3 and 4), whereas injections in the forelimb representation resulted in label further detached from this border (their case 7). In addition, injections in the representation of the facial musculature labeled neurons in even more rostral locations within 6M, but only in the lateral half of this area. These observations are compatible with the proposed somatotopic gradient found across the macaque SMA and pre-SMA (Luppino et al. 1993). Interestingly, in light of the present results, projections from rostromedial 6M to M1 were very sparse (Burman et al. 2014a).

In summary, whereas the information on connections is highly compatible with the proposed somatotopic organization of the SMA and pre-SMA in the macaque, it may further suggest that the border between these subdivisions may not be perfectly perpendicular to the rostrocaudal axis. This hypothesis, illustrated in Figure 11, could be one explanation for the observation that afferents from the facial representations of M1 and somatosensory areas (3a, 3b, 1/2) were relatively sparse in the present materials (as none of the injections was located in rostrolateral 6M). However, it must also be considered that studies in New World monkey species including marmoset, owl and squirrel monkey have typically reported connections between the SMA to the lateral parts of M1 and somatosensory areas as being weak, sparse or inconsistent (Krubitzer and Kaas 1990; Guldin et al. 1992; Stepniewska et al. 1993; Huffman and Krubitzer 2001; Burman et al. 2014a).

**Figure 11:**
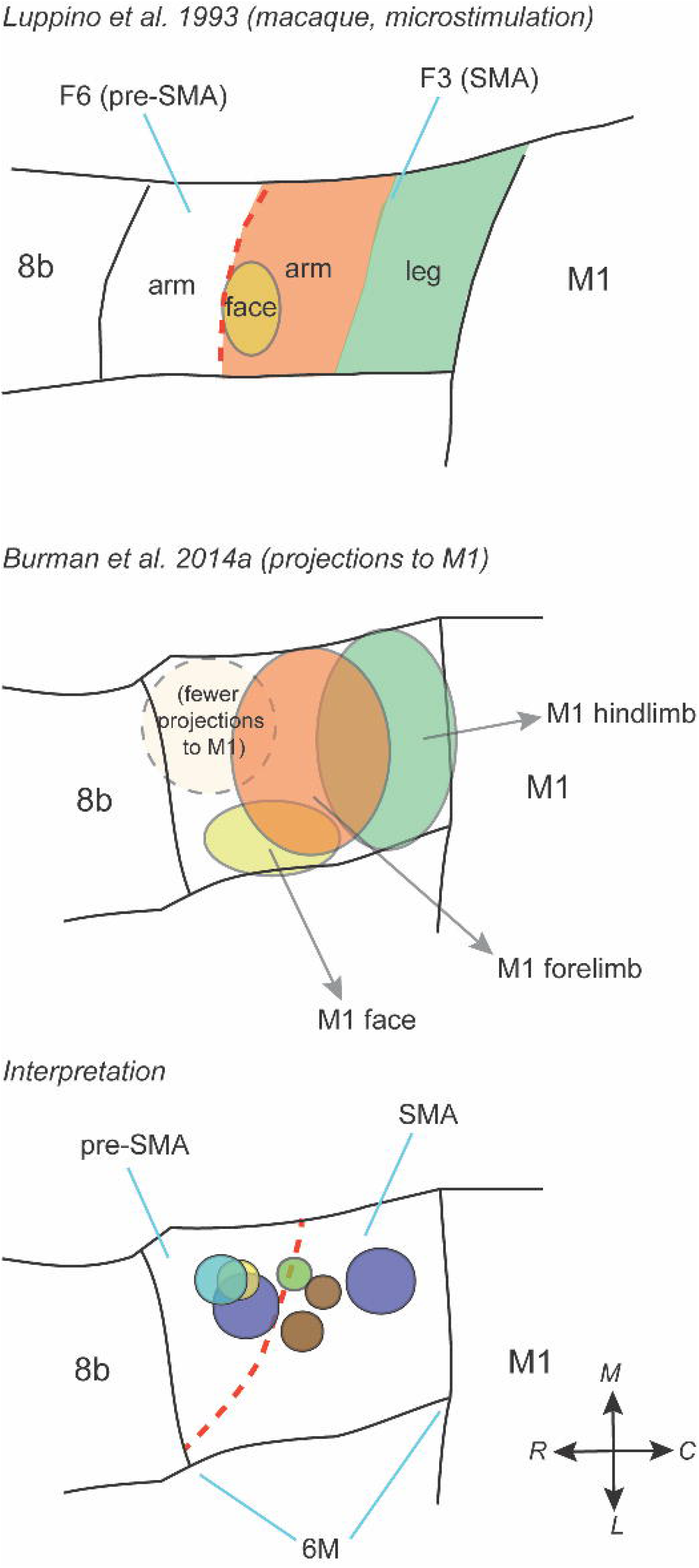
Comparison of the topographic organization of the macaque and marmoset medial area 6. **A:** Organization in the macaque, based on microstimulation results summarized by Luppino et al. (1993). To provide a ready comparsion with the marmoset data, the figure illustrates a left hemisphere, with the midline dorsally. **B:** Summary of the connectional data from Burman et al. 2014a (retrograde tracer injections in M1). **C:** Interpretation of the location of the present injections relative to the somatotopy shown in B. The orientation of the maps is shown by the insert on the bottom right (R-rostral, C-caudal, M-medial, L-lateral).

### Organization of sensorimotor pathways in the marmoset

Present and past tracing results delineate multiple parallel pathways through the marmoset motor and premotor areas, and highlight the likelihood of different contributions to movement control (for evidence in other species, see Gharbawie et al. 2011; Caminiti et al. 2017). For example, compared to the adjacent M1 and 6DC, 6M receives more numerous projections from prefrontal areas and rostral midline areas. On the other hand, 6M receives on average fewer inputs (by about one quarter) from the dorsorostral posterior parietal cortex and only weak inputs from the somatosensory areas. Assuming that connectional strength signifies functional specificity, 6M appears to be less influenced by sensory and spatial information, in comparison with both M1 and 6DC. These observations are consistent with the relative lack of responsiveness during somatic peripheral stimulation in macaques (Brinkman and Porter 1979; Wise and Tanji 1981) and lend support to a specialized role in mediating self-paced and temporal sequences of movements (Tanji 2001).

The rostral midline input to area 6M originates not only in areas 24c and 24d, which have been regarded as homologues of the cingulate motor areas (present results and Bakola et al. 2015), but also anterior cingulate association areas 24b and (to a lesser extent) 24a. Indeed, the present experiments provide the first evidence regarding the connections of area 24d, in doing so confirming its identity as a premotor field. In terms of location relative to M1 and 6M, marmoset area 24d appears to closely correspond to the macaque’s area CMAd (He et al. 1995). Based on a single injection centered in this area, our results reveal all of the connections reported in the macaque (Hatanaka et al. 2003), but suggest a more extensive network including somatosensory, posterior parietal and prefrontal inputs to 24d. Relative to other motor fields, area 24d differs by the increased emphasis on projections from posterior cingulate areas, in particular areas 23a and 23b.

One notable aspect of the parietal input to 6M is that most projections stem from dorsal posterior parietal areas PE, PEC and 31. As in the macaque, parietal connections also originate in smaller proportions in inferior parietal areas PF and PFG. However, studies in this species also describe connections with the medial intraparietal area, MIP (Petrides and Pandya 1984; Luppino et al. 1993; Morecraft et al. 2012; Bakola et al. 2017), which were not evident in the present study. We have previously seen that 6DC in the marmoset also receives largely restricted parietal projections (mainly from PE; Burman et al. 2014b). Our past and present results in the marmoset can be tentatively interpreted within the context of differential cortical mechanisms for movement planning in a less dexterous primate, but the issue requires further investigation. In particular, the precise extent of area MIP in the marmoset has only been proposed on the basis of location and cytoarchitecture (Paxinos et al. 2012), but not confirmed physiologically.

Relative to the dorsorostral premotor cortex (6DR, Burman et al. 2014a), 6M receives much sparser projections from the posterior midline and caudal parietal visuomotor areas (e.g. areas 23, areas LIP and PGm). Areas 6M and 6DR also differ in terms of their prefrontal connectivity, with 6DR receiving more robust projections from subdivisions of area 8 that host the marmoset frontal eye fields (Burman et al. 2006; Reser et al. 2013; Ghahremani et al. 2017; Feizpour et al. 2021). The distinctiveness of 6M and 6DR in the marmoset, judged by anatomical data, parallels descriptions in the macaque (Luppino et al. 1993; Luppino et al. 2003), and suggests that in both species 6DR is more involved in abstract levels of motor planning, as well as in visual and oculomotor processes. Further, studies in macaques (Schlag and Schlag-Rey 1987; Russo and Bruce 2000; Fujii et al. 2002; Moschovakis et al. 2004; Amiez and Petrides 2009), owl monkeys (Gould et al. 1986; Stepniewska et al. 1993) and capuchin monkeys (Tian and Lynch 1996) have reported eye-movement activity in rostral premotor cortex medial to the classic frontal eye field. The more dorsal locations from which eye movements can be evoked with low-threshold currents are commonly referred to as the supplementary eye field, but relationships with architectonic fields remain a matter of contention. Our tracing data suggest that in the marmoset area 6M is not a likely an eye-related region, in line with evidence from anatomical (Huerta and Kaas 1990) and microstimulation studies (Luppino et al. 1991; Matsuzaka et al. 1992; Fujii et al. 2002; Yang et al. 2008) in the macaque. Rather, they suggest that the supplementary eye field may exist within marmoset area 6DR, where visual receptive fields have recently been observed (Feizpour et al. 2021).

The sparse direct projections from the ventral premotor cortex to marmoset area 6M contrast the generally denser connections reported in macaques (Luppino et al. 1993; Morecraft and Van Hoesen 1993; Ghosh and Gattera 1995) and capuchin monkeys (Dum and Strick 2005); note, however, that other studies have reported moderate to weak connections (Gerbella et al. 2011; Morecraft et al. 2012). A reciprocal, relatively weak pathway has been described after injections in the marmoset area 6Va (Burman et al. 2015). The strength of this connection appears to depend on somatic representation, with stronger connections targeting the forelimb area, Luppino 1993). Ventral parts of the premotor cortex are essential for the visuomotor control of grasping actions, but relevant contributions from medial subdivisions of area 6 have been the focus of only a few studies (Lanzilotto et al. 2016).

### Medial premotor-prefrontal pathways

We found a substantial projection from the granular frontal cortex to the rostral part of area 6M, similar to studies of the macaque preSMA/F6 (Luppino et al. 1993; Morecraft et al. 2012). The bulk of these prefrontal afferents originated in area 8b. Afferents to the macaque medial premotor cortex also arrive from a corresponding brain location (i.e., immediately rostral to the preSMA) even though area 8b has not been mapped consistently in that species (e.g., Luppino et al. 1993). Macaque studies have described additional projections from the dorsolateral prefrontal region around the principal sulcus (areas 9 and 46), but these were either absent or very sparse in our data. However, other studies have reported that connections between the medial premotor cortex and the periprincipal cortex are very weak (Bates and Goldman-Rakic 1993; Takada et al. 2004). It has been suggested that the connections between the periprincipal region and the preSMA/F6 are related to motor programs for reaching and grasping (Lu et al. 1994; Borra et al. 2019). Thus, another possibility is that the paucity of projections from areas 9/46 to rostral 6M is a further reflection of the limited manual skills of marmosets. Intriguingly, sparse connections from a small region near the rostral end of orbitofrontal area 13L were also observed in 5 of the injections in area 6M, the significance of which remains unclear.

Another aspect of prefrontal-premotor connectivity, which has been relatively overlooked, relates to the distributed networks for auditory and language processing. Based on MRI studies in humans, the medial premotor cortex has been suggested as a site for auditory integration, even when no overt motor response is required (Lima et al. 2016). A similar role for the non-human primate medial area 6 has not been reported, and any links with the auditory system must be indirect. Such indirect route would likely be through area 8b, which has been proposed as part of the network of areas for recognition of complex auditory stimuli and for orienting ear and eye movements (Luchetti et al. 2008; Lanzilotto et al. 2013). Human tractography and dissection studies (Catani et al. 2012; Vergani et al. 2014) have also suggested the existence of axonal pathways between the inferior frontal cortex and the medial part of area 6, which were proposed to serve linguistic processing. Some evidence of such pathways exists in the macaque, suggesting an evolutionary predecessor (Petrides and Pandya 2002; Gerbella et al. 2010), and the SMA has been implicated in learning sound sequences which could be used for communication (Archakov et al. 2020). In marmosets, projections from ventrolateral prefrontal areas (45/ 47) to area 6M were only obvious following 2 injections (CJ163-FB and CJ164-CTBg), which were near the rostral limit of this area. Projections from the ventral premotor cortex (areas 6Va/ 6Vb), which have been implicated in the production and perception of vocalizations (Coudé et al. 2011), provide another potential pathway to mediate the involvement of area 6M in vocal communication.

As part of the present study we had the opportunity to re-examine the connections of area 8b, which had been initially examined by Reser et al. (2013) with the aim of assessing differences between cytoarchitectural subdivisions of area 8. The main conclusions of the previous study were supported: area 8b is characterized by a wide and varied range of connections, including the entire forntal lobe, anterior and posterior cingulate areas, and high-order sensory and sensory integration areas located in dorsal temporal and ventral parietal cortex. This variability is unlikely to be explainable from the fact that 2 of the injections did not extend to the infragranular layers (Table 1), as it consisted of different combinations of labeled areas across injection sites, rather than lack of some projections in these cases (8 and 11) relative to those extending to layer 6 (cases 9 and 10). Among premotor areas, its main connections are formed with areas 6DR and 6M, although the present extended sample revealed that some parts of area 8b also receive projections from the ventral premotor cortex (Fig. 9E). These results hint at a role of area 8b in integration of sensory information in preparation to attention shifts and facilitation of motor responses to specific aspects of the extrapersonal world (e.g. Lanzilotto et al. 2013).

## Conclusion

Our results reveal the fundamental similarity of the premotor cortex near the midline across primates, despite motor behaviour patterns that vary from vertical clinging and leaping in marmosets, to the highly dexterous manipulation by humans and macaques. In all of these species the connections of area 6M revealed a dominance of afferents from cortical areas associated with motor control, pointing to the medial premotor cortex as a critical node in the neural circuitry for action. By providing quantitative comparisons with adjacent areas, our data reveal the uniqueness of area 6M’s connectivity pattern, and provide the basis for models that can incorporate anatomical information to the growing body of physiological data (e.g. Ebina et al. 2018, 2019; Tia et al. 2021) in this species. The evidence for rostral and caudal subdivisions (putative homologues of the SMA and pre-SMA) adds to other data (e.g. Majka et al. 2019) which, in the future, may prompt the preparation of new stereotaxic atlases which include finer parcellations of the marmoset cortex, within currently recognized cytoarchitectural areas.

## Acknowledgements

Funded by grants from the Australian Research Council (DE120102883, DP140101968, CE140100007), National Health and Medical Research Council (1020839, 1082144, 1194206), European Research Council (FP7-PEOPLE-2011-IOF 300452; H2020-MSCA-734227–PLATYPUS), and the National Science Centre (2019/35/D/NZ4/03031). The authors would like to thank Daria Malamanova, Karyn E. Richardson, and Kirsty Watkins for contributions related to histology and microscopy, Cecilia Cranfield for helping edit the text and figures, and Dr. David H. Reser for constructive comments at many stages of this project. Slide scanning was performed by the Monash Histology Platform.

