## Supplementary Table 1 for "Afferent connections of cytoarchitectural area 6M and surrounding cortex in the marmoset: putative homologues of the supplementary and pre-supplementary motor areas"

**Supplementary Table 1: Abbreviations of names of cortical areas**

1-2	areas 1 and 2 of cortex

10	area 10 of cortex

11	area 11 of cortex

13L	area 13 of cortex lateral part

13M	area 13 of cortex medial part

13a	area 13a of cortex

13b	area 13b of cortex

14C	area 14 of cortex caudal part

14R	area 14 of cortex rostral part

23V	area 23 of cortex ventral part

23a	area 23a of cortex

23b	area 23b of cortex

23c	area 23c of cortex

24a	area 24a of cortex

24b	area 24b of cortex

24c	area 24c of cortex

24d	area 24d of cortex

25	area 25 of cortex

29a-c	area 29a-c of cortex

29d	area 29d of cortex

30	area 30 of cortex

31	area 31 of cortex

32	area 32 of cortex

32V	area 32 of cortex ventral part

35	area 35 of cortex

36	area 36 of cortex

3a	area 3a of cortex (somatosensory)

3b	area 3b of cortex (somatosensory)

45	area 45 of cortex

46D	area 46 of cortex dorsal part

46V	area 46 of cortex ventral part

47L	area 47 (old 12) of cortex lateral part

47M	area 47 (old 12) of cortex medial part

47O	area 47 (old 12) of cortex orbital part

4ab	area 4 of cortex parts a and b (primary motor, M1)

4c	area 4 of cortex part c (primary motor, M1)

6DC	area 6 of cortex dorsocaudal part

6DR	area 6 of cortex dorsorostral part

6M	area 6 of cortex medial (supplementary motor) part

6Va	area 6 of cortex ventral part a

6Vb	area 6 of cortex ventral part b

8C	area 8 of cortex caudal part

8aD	area 8a of cortex dorsal part

8aV	area 8a of cortex ventral par

8b	area 8b of cortex

9	area 9 of cortex

AI	agranular insular cortex

AIP	anterior intraparietal area of cortex

APir	amygdalopiriform transition area

CL	auditory cortex caudolateral area

CPB	auditory cortex caudal parabelt area

DI	dysgranular insular cortex

Ent	entorhinal cortex

FST	fundus of superior temporal sulcus area of cortex

GI	granular insular cortex

Gu	gustatory cortex

IPro	insular proisocortex

LIP	lateral intraparietal area of cortex

M1	primary motor area (cytoarchitectural areas 4ab and 4c)

ML	auditory cortex middle lateral area

MST	medial superior temporal area of cortex

OPAl	orbital periallocortex

OPro	orbital proisocortex

OPt	occipito-parietal transitional area of cortex

PE	parietal area PE

PEC	parietal area PE caudal part

PF	parietal area PF (cortex)

PFG	parietal area PFG (cortex)

PG	parietal area PG

PGM	parietal area PG medial part (cortex)

PGa/IPa	Area PGa andIPa (fundus of superior temporal ventral area)

Pir	piriform cortex

ProM	proisocortical motor region (precentral opercular cortex)

ProSt	prostriate area

RPB	auditory cortex rostral parabelt

S2	secondary somatosensory cortex

S2PR	secondary somatosensory cortex parietal rostral area

S2PV	secondary somatosensory cortex parietal ventral area

STR	superior temporal rostral area (cortex)

TE1	temporal area TE1 (inferior temporal cortex)

TE2	temporal area TE2 (inferior temporal cortex)

TE3	temporal area TE3 (inferior temporal cortex)

TEO	temporal area TE occipital part

TF	temporal area TF

TFO	temporal area TF occipital part

TH	temporal area TH

TL	temporal area TL

TLO	temporal area TL occipital part

TPO	temporo-parieto-occipital association area (superior temporal polysensory)

TPPro	temporopolar proisocortex

TPt	temporoparietal transitional area

V6a	visual area 6a (posterior parietal medial area)

VIP	ventral intraparietal area of cortex
