## Supplementary figures and images for "Afferent connections of cytoarchitectural area 6M and surrounding cortex in the marmoset: putative homologues of the supplementary and pre-supplementary motor areas"

### Supplementary Figure S1

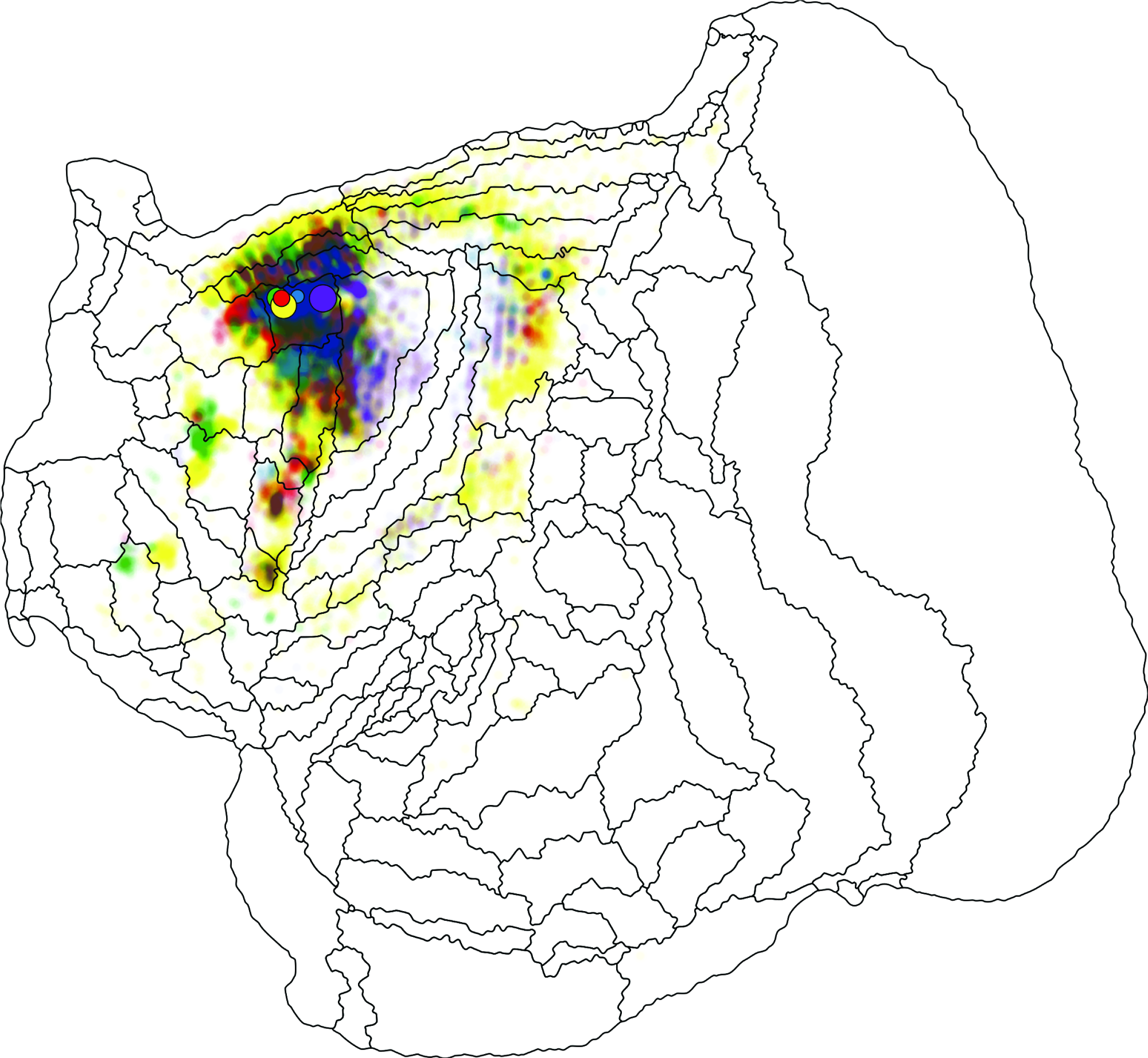
